# Depletion of membrane cholesterol modifies structure, dynamic and activation of Na_v_1.7

**DOI:** 10.1101/2024.02.21.581348

**Authors:** Simone Albani, Vishal Sudha Bhagavath Eswaran, Alessia Piergentili, Paulo Cesar Telles de Souza, Angelika Lampert, Giulia Rossetti

## Abstract

Cholesterol is a major component of plasma membranes and unsurprisingly plays a significant role in actively regulating the functioning of several membrane proteins in humans. Notably, recent studies have shown that cholesterol depletion can also impact transmission of potentially painful signals in the context of peripheral inflammation, via hyperexcitability of the voltage-gated sodium channel (Na_v_) subtype 1.9, but the structural mechanisms underlying this regulation remain to be elucidated. In this study, we focus on the role of cholesterol depletion on Na_v_1.7, which is primarily expressed in the peripheral sensory neurons and linked to various chronic inherited pain syndromes. Coarse-grained molecular dynamics simulations shed light on the dynamic changes of the geometry of Na_v_1.7 upon membrane cholesterol depletion: A loss of rigidity at key structural motifs linked to activation and fast-inactivation is observed, as well as changes in the geometry of drug-binding regions in the channel. Loss of rigidity in cholesterol depleted conditions should allow the channel to transition between different gating states more easily. *In-vitro* whole-cell patch clamp experiments on HEK293t cells expressing Na_v_1.7 validated these predictions made *in silico* at the functional level. Hyperpolarizing shifts in the voltage-dependence of activation and fast-inactivation were observed along with an acceleration of the time to peak and onset kinetics of fast inactivation. These results underline the critical role of membrane composition, and of cholesterol in particular, in influencing Na_v_1.7 gating characteristics. Furthermore, our results hint to a key role of the membrane environment in affecting drug effects and in pathophysiological dysregulation, sharpening our approaches for analgesics design.

**Supplementary data:** https://doi.org/10.5281/zenodo.10829175

## Introduction

Cholesterol is a major factor in influencing the biophysical properties of plasma membranes, where it constitutes 30–45% of lipid molecules^1,2^. Several studies have shown that cholesterol modulates the function of a wide range of receptors and ion channels embedded in the membrane via specific, i.e., direct, ligand-like interactions, and non-specific indirect mechanisms^3–5^. For instance, cholesterol impacts the biophysical, microstructure and mechanical properties of the membrane, in turn impacting on the trafficking and expression level of several proteins (for a review see ref.^6^). Also, the molecular rearrangements resulting in the gating of ion channels, the transporting processes and the conformational transitions of transmembrane receptors are mediated by the surrounding lipid bilayer. Thus, all of these functions can be modified by cholesterol^6^.

Recently, unexpected functions of cholesterol in controlling transmission of potentially painful (nociceptive) signals were uncovered, involving voltage-gated sodium channels (Na_v_s). In the human body, nine different subtypes of Na_v_s are expressed (Na_v_1.1 to Na_v_1.9), which are responsible for the fast upstroke of the action potential. They open when the membrane potential in their vicinity depolarizes, allowing sodium ions to flow into the cell^7^. Na_v_1.7, Na_v_1.8 and Na_v_1.9, are expressed predominantly in the peripheral nervous system and are involved in the transduction and propagation of nociceptive signals^8^. A recent study suggests that pharmacological depletion of cellular cholesterol entails sensitization of nociceptive neurons potentially mediated via Na_v_1.9, promoting mechanical and thermal hyperalgesia^9^. A similar effect was observed with inflammatory mediators, which were reported to support the partitioning of Na_v_1.9 channels from cholesterol-rich microdomains of the plasma membrane (lipid rafts), in which also sphingolipids and gangliosides are accumulated^10^, to cholesterol-poor non-raft regions of the membrane. Low-cholesterol environment enhances voltage-dependent activation of Na_v_1.9 channels leading to enhanced neuronal excitability^9^. Na_v_1.8 was also shown to activate more easily in cholesterol-depletion conditions, when the channel is disassociated from lipid rafts^11^, suggesting that cholesterol can be a powerful modulator of nociception via Na_v_s. Although several cholesterol interaction sites on Na_v_1.9 were suggested, the molecular understanding of cholesterol-regulated hyperexcitability is not yet known^9^.

In this study, we employed a range of computational and experimental techniques to assess the transferability of cholesterol-dependent effects to the subtype Na_v_1.7 (**Fig. 1**), which was shown to be involved in human pain perception^12,13^.

**Figure 1.**
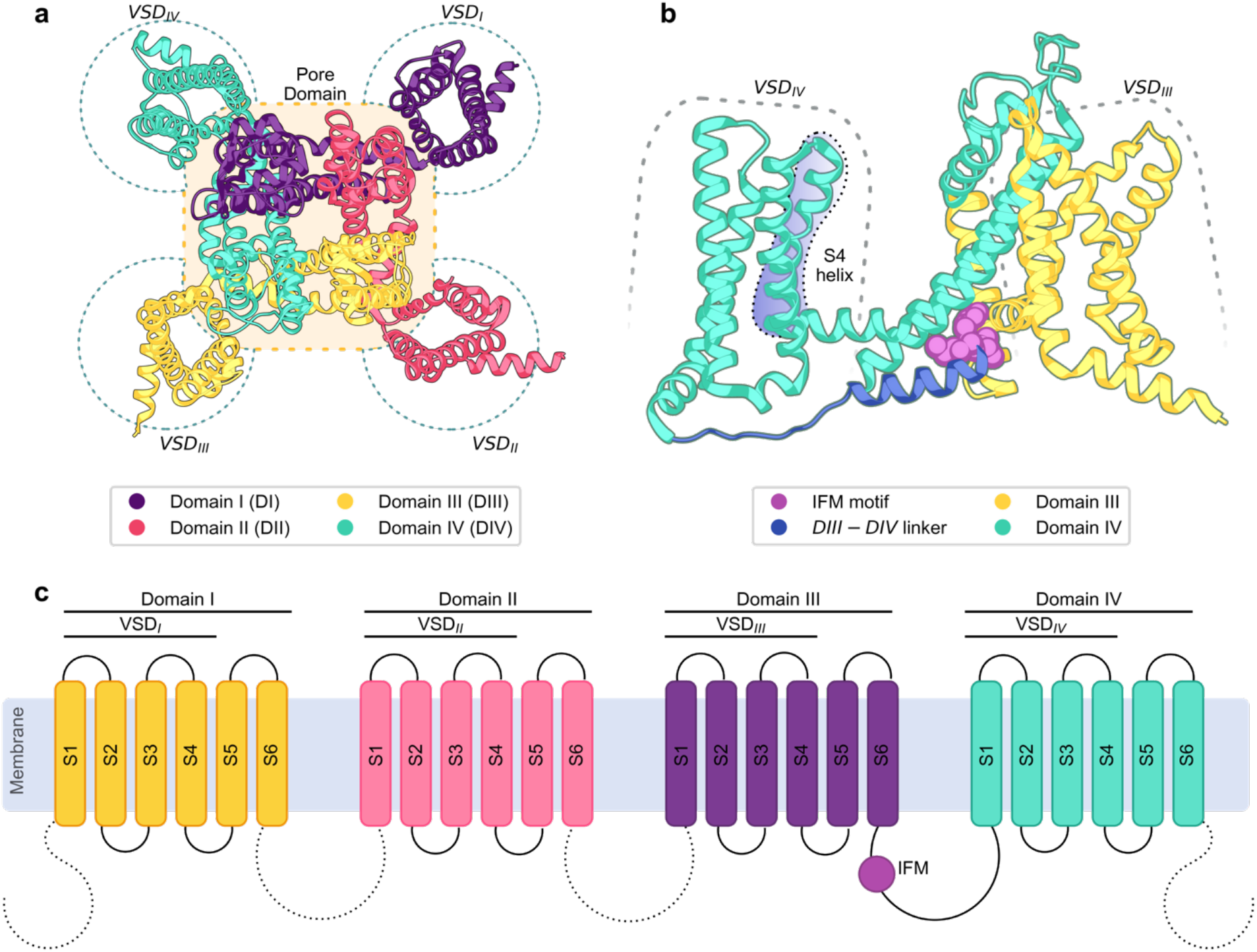
Structural overview of Na_v_1.7 channel. **a**. Extracellular view of the α subunit of Na_v_1.7 channel. It is a single-chain protein composed of four domains, each containing 6 transmembrane (TM) helices named from S1 to S6. In each domain, helices S1 to S4 are part of the voltage-sensing domain (VSD), while the helices S5 and S6 are part of the Pore Domain (PD)^14^. **b**. Side view of Domains III and IV. The fourth transmembrane helix (S4) in each domain, due to its positive residues, moves towards the extracellular side when the membrane is depolarized and is the main driver for voltage sensing. When Na_v_ channels open and conduct sodium ions, the ion flow is interrupted after a few milliseconds by the process called “fast inactivation” ^15^. During this process, a hydrophobic group of residues (Ile-Phe-Met, called IFM motif or inactivation particle) in the linker between Domains III and IV binds to a corresponding pocket in between the VSD of DIII and the PD, and allosterically restricts the pore^16,17^. **c**. Schematic illustration of the simulated α subunit of Nav1.7, showcasing the TM helices, the DIII-DIV linker, the IFM motif, and the absent (dashed lines) terminal domains and inter-domain linkers.

Not only do we provide evidence that cholesterol depletion causes hyperexcitability of Na_v_1.7, similar to what was observed for Na_v_1.9 and Na_v_1.8, we also provide a molecular interpretation of how cholesterol interaction with Na_v_1.7 structure impacts the dynamic features of the channel^9,11^. Notably, such structural and dynamic changes significantly impact Na_v_1.7 interaction sites of known drugs, suggesting an effect on channel druggability in the presence of membrane-lipid modifiers such as inflammatory agents.

## Results

We conducted coarse-grained molecular dynamics simulations to investigate the effect of membrane composition on the geometry of the α subunit of the human Na_v_1.7 channel in a fast-inactivated state (**Fig. 1**). Specifically, we simulated the channel in three different membrane compositions: The first condition was designed to mimic a lipid raft in a neuronal membrane with a complex composition of lipids; these includes cholesterol and the ganglioside GM1 as main components (see methods); the second condition was composed solely of phosphatidylcholine (POPC, see methods for further details) to mimic a full detachment from the lipid raft; the third condition has the same composition of the first one but the cholesterol was depleted and replaced with phosphatidylcholine. This last membrane was designed to isolate the effect of cholesterol, if any, from the one of the other components. Experimentally, we performed whole-cell voltage clamp experiments in HEK293t cells expressing Na_v_1.7 under control conditions and during pharmacologically induced cholesterol depletion.

### Properties of the membrane and specificity of lipids interaction

The lipid raft-like membrane displayed a smaller area-per-lipid (APL 0.506 ± 0.003 nm^2^, **Fig. 2a**) and an increased thickness (38.98 ± 0.25 Å, **Fig. 2b**) compared to the cholesterol-depleted (APL 0.642 ± 0.004 nm^2^, and thickness 37.54 ± 0.26 Å) and the single-component membrane (APL 0.661 ± 0.004 nm^2^, and thickness 38.59 ± 0.21 Å). A tighter packing of the lipids (here expressed as APL) was expected in the membrane containing cholesterol^18^. The reduced thickness in the cholesterol-depleted membrane is also in line with previous computational and experimental studies carried out with simple membrane compositions in different cholesterol concentrations^19–21^. Despite these morphological changes, no significant variation in membrane curvature was observed (gaussian curvature in the range of −0.0125 to 0.008 Å^-2^). As a next step, we tested for potential direct interactions between the membrane and key structural elements of Na_v_1.7. Specifically, we calculated the occupancy of each membrane component around the channel to determine the high-occupancy regions (i.e., regions where a given component is more frequently present) (**Fig. S1** and **S2**). In the single-component membrane, no specific interactions between the protein and the phosphatidylcholine molecules were identified. This is not the case for the components of other two membrane types: Cholesterol and Ganglioside molecules emerged as able to establish localized interactions with the channel. Such identified high-occupancy regions were compared with existing knowledge and pertinent literature regarding known interaction sites of Na_v_ channels at the level of the membrane.

**Figure 2.**
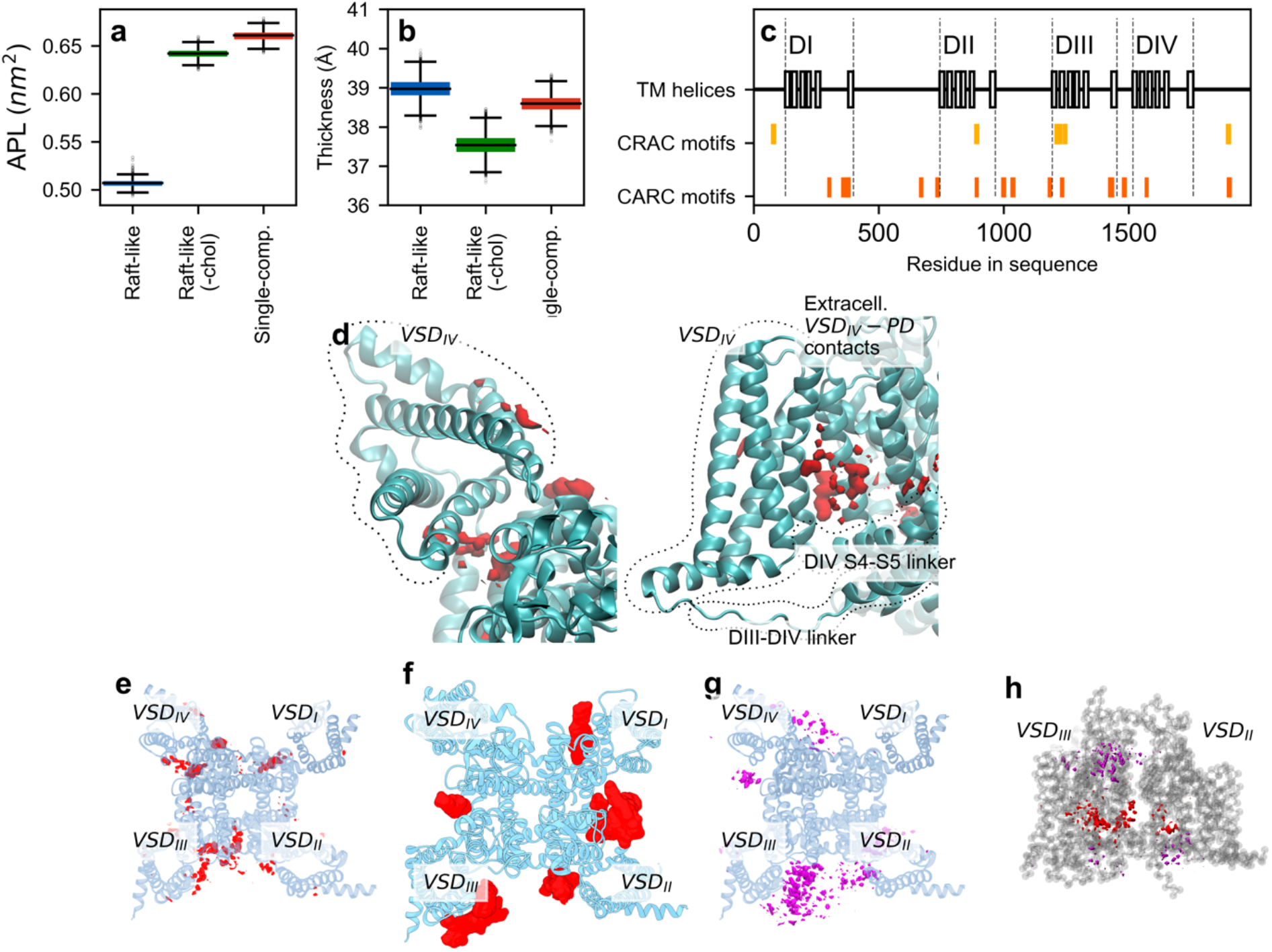
Analysis of membrane properties and lipid-protein interactions. **a**. Area per lipid (APL) values calculated over 12 µs per each membrane type (1 sample/ns). APL is considered a measure of how tightly packed the lipid molecules are within the membrane. **b**. Membrane thickness, measured as the average distance of phosphates groups between the two leaflets and calculated over 12 µs per each membrane type (1 sample/ns). **c**. Sequence of CRAC and CARC motifs along protein sequence. The closest correspondences between CARC/CRAC motifs and TM helices are the partial overlap of two CRAC motifs with the end of DIII S1 (overlap on residues 1209-1211 of isoform 1) and DIII S2 (on residues 1242-1243), and a CARC motif in the middle of DIII-S2 (on residues 1231-1238) (Fig. 2c) **d**. Detailed extracellular (left) and side (right) view of cholesterol occupancy around VSD_IV_ (red isosurface) **e**. Extracellular view of cholesterol occupancy around the overall Na_v_1.7 structure (red isosurface). **f**. Cholesterol and cholesteryl hemisuccinate molecules positioning around eukaryotic Na_v_ as previously observed in CryoEM experimentsPDB IDs: 6NT3,6NT4, 7W9K, 7W9L, 7W9P, 7W9T, 7XVE, 7XVF) (red surface), overimposed onto extracellular view of Na_v_1.7^15,24,25^. Cholesteryl hemisuccinate is not naturally occurring in cells^26^, but it can be used as a detergent and stabilizer of membrane proteins^27^, and as such it is often found interacting with eukaryotic Nav channels. The common placements between CryoEM and simulations are marked here with roman numerals (i to iv), see main text. **g**. Extracellular view of ganglioside GM1 occupancy around Na_v_1.7 (purple isosurface). **h**. Cholesterol (red) and GM1 ganglioside (purple) occupancy, visualized from the side of the DIII-DIV fenestration. The fenestration is formed by S5 and S6 of DIII and DIV, and situated between VSD_II_ and VSD_III_.

For cholesterol, we first compared the binding sites suggested for Na_v_1.9, as these motifs are highly conserved in Na_v_1.7 (**Table 1**). Specifically, we looked for so-called “CRAC” and “CARC’’ motifs, i.e., particular sequences, that when positioned near the membrane-solvent boundary like in Na_v_1.9, can serve as reliable indicators for enhanced cholesterol binding affinity^22^. However, these motifs, when present in Na_v_1.7, either do not fall in the TM regions, or, when they do, they do not align to the lipid bilayers. This suggests that such motifs do not constitute good indicators for cholesterol binding in Na_v_1.7. Therefore, we next compared high cholesterol occupancy regions (at least 4% of the sampled frames, in 1 Å^3^ voxels) in our simulations with the positioning of cholesterol and its derivatives in structures obtained by CryoEM (**Fig. 2de**). Four common placements between simulations and CryoEM experiments were found: (i) above the S4-S5 linker of DI; (ii) on the surface of VSD_III_, close to S3; (iii) at the entrance of the fenestration between VSD_III_ and VSD_IV_; (iv) in proximity of VSD_II_ S1 (**Fig. 2ef**). However, in the last two cases the binding appears to be less stable as the occupancy is lower than 7%. Two additional binding sites can be observed in the simulations hovering the S4-S5 linker of the domains DIII and DIV (**Fig. 2de**). In particular, S5 of DI interacts frequently with cholesterol on both of its sides exposed to the membrane, thus rendering the mechanical hinge created by the S4-S5 linker of DIV and non-bonded interactions between VSD_IV_ and S5 and S6 of DI particularly stabilized (**Fig. 2d**).

**Table 1:**
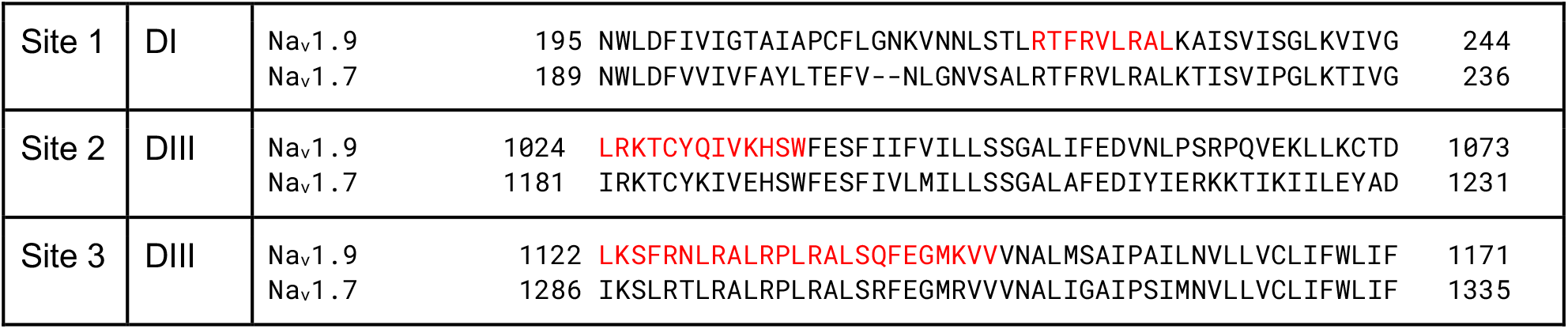
Pairwise alignment. Pairwise alignment of mouse Na_v_1.9 (uniprot ID: Q9R053) and human Na_v_1.7 (uniprot ID: Q15858-1) sequences in correspondence of the cholesterol binding sites found by Amsalem et al. (highlighted in red)^9^.

Even though ganglioside molecules, too, have been demonstrated to increase the excitability of Na_v_ channels^25^, there are no known binding sites for these compounds.

In our lipid raft-like simulations we found that ganglioside molecules predominantly occupy the membrane-solvent interface in proximity of the VSDs (**Fig. 2g**). Notably, the highest occupancy is observed in the vicinity of VSD_III_, where ganglioside molecules complement the cholesterol densities (**Fig. 2h**). In the cholesterol-depleted membrane gangliosides exhibit interactions with the protein in the same regions, but an additional concentration of gangliosides is observed at the entrance of the DI-DII fenestration. Intriguingly, in all the simulations with raft-like membranes, VSD_IV_ encounters multiple ganglioside clusters close to the S4-S5 linker, potentially constraining its range of motion (**Fig. 2g** and **S2**).

### Membrane composition shapes 4-fold symmetry of Na_v_1.7 *in silico* modeling

Asymmetric rearrangements in the PD have been associated with channel opening and closing. Specifically, monitoring the position of four residues located in the pore at the level of the inactivation gate enables the observation of a shift in the shape of the pore from a square to a rhomboid during the transition from a closed to an open or inactivated state in the rat Na_v_1.5 channel^17^. We first evaluated whether fluctuations of the square shape of the channel occur naturally, within a timescale shorter than that required for VSD activation or channel opening. To achieve this aim, we measured changes in the diagonals of such a square in all three simulated environments, specifically, the angle between vectors connecting VSD_I_ to VSD_III_ and VSD_II_ to VSD_IV_(**Fig. 3a** and **S3**). Among the deposited human Na_v_1.7 structures, values of such angles are around 84.5° (84.45°, 84.38°, 84.65° for structures with PDB IDs 6J8G, 7W9P, 8G1A respectively). Such angle is fairly conserved in our simulations of lipid raft-like membranes (86.64 ± 1.48° and 84.12 ± 2.03° with and without cholesterol, respectively). This is not the case for the single-component membrane, where the angle values broaden up, assuming a bimodal distribution with peaks at 80.78° and 90.23° (**Fig. 3b**), suggesting a geometrical distortion.

**Figure 3.**
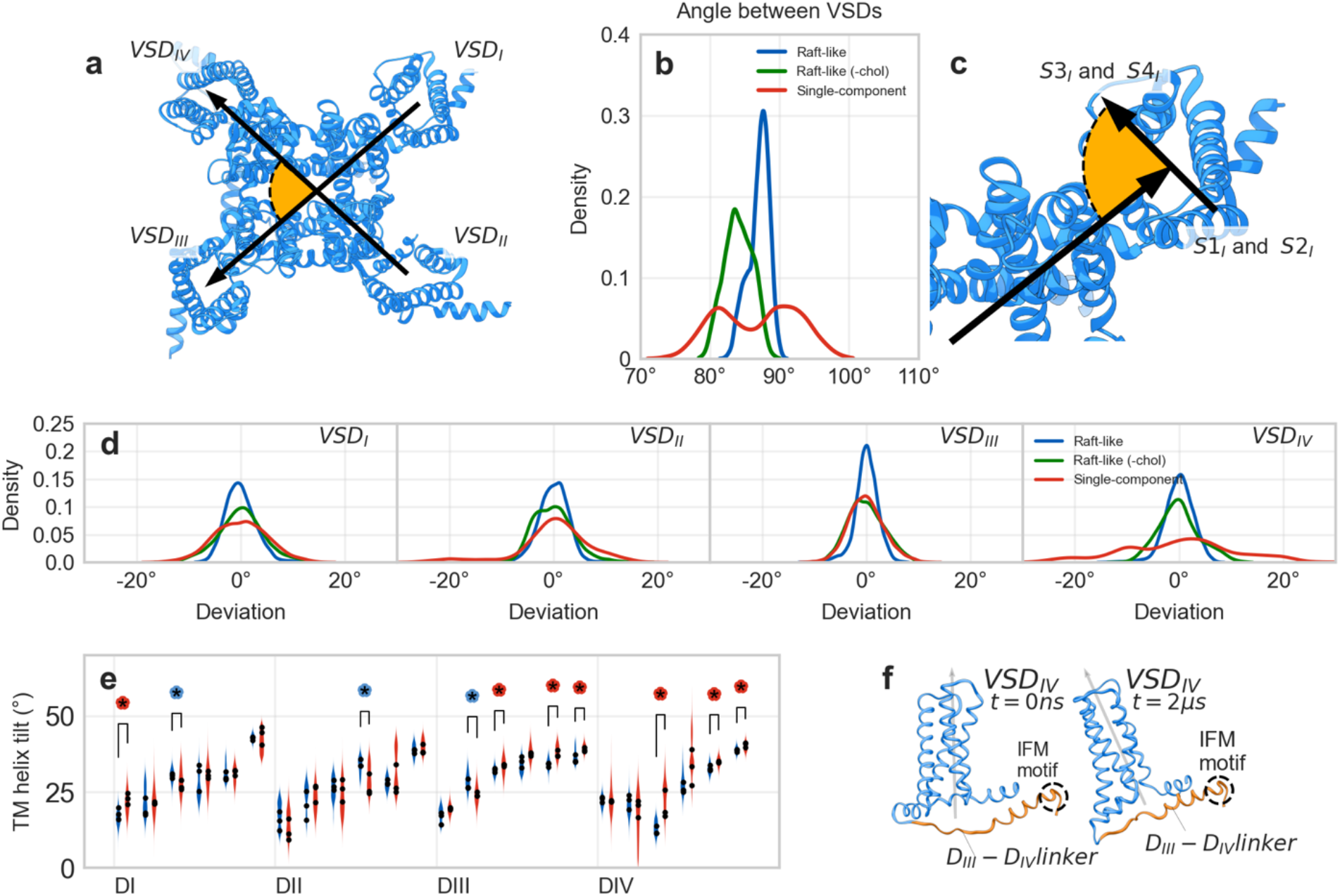
Structural changes in VSDs. **a**. Extracellular view of Na_v_1.7 structure and visualization of the selected angle to monitor 4-fold symmetry of the channel. **b**. Distribution of values of the angle between opposite VSDs for the three membrane compositions. **c**. Extracellular view of VSD_I_ and definition of the angle to monitor VSD orientation with respect to the pore domain (see Figure S4 for further details). **d**. Distribution of angles of rotation of the four VSDs for the three membrane compositions. 0° corresponds to the average orientation of the VSD in the replica. For panel **b** and **d** the following color scheme is implemented: blue, green and red for cholesterol-rich lipid raft-like, cholesterol-depleted lipid raft-like and single-component membranes, respectively. **e**. Distribution of tilt angle for the 24 TMs in the lipid raft-like and single-component membrane simulations. The average of each replica per each membrane is represented by black circles. A coloured stroked asterisk marks a significant difference in the tilt of an TM between the two conditions (see test in methods). Asterisks are color-coded as in panel **b** and **d. f**. The tilt of VSD_IV_ in the second replica of single-component membrane simulations.

### Rotation of VSDs and changes in PD-VSD interface

CryoEM experiments show that the translocation in space of the VSD around PD is accompanied by a rotation of the VSD around its vertical axis, which in turn alters the interaction pattern between VSD and PD^25,28^. This observation could be linked to a process of deactivation of the VSD, as seen in a recent Na_v_1.7 CryoEM structure. In this study, mutagenesis-induced deactivation of VSD_I_ resulted in the domain tracing an angle of approximately 18° around the PD, oriented towards VSD_IV_^25^. Alternatively, a more extensive sampling may reveal similar phenomena, as demonstrated in the activated VSD_I_ of a wild-type Na_v_1.8, which was captured after a rotation of 12° around the core of the protein^28^. Although this protocol cannot simulate the rearrangements caused by a VSD deactivation, we observed this rotation in all four domains across our simulations and found that it is in the majority of the frames limited to a range of 6° and normally distributed. Some relevant exceptions appear when the channel is simulated in the single-component membrane (**Fig. 3cd** and **S5**): The distribution of angles for VSD_II_ is wide (min.: −23.4°, max.: +32.4°, st.dev.: 10.8° with respect to t=0 μs) and skewed towards positive values, due to a slightly more favorable, albeit still infrequent, placement of this DI in proximity of VSD_III_ in one of the replicas (**Fig. 3d**, and **S5**); Rotations of the VSD_IV_ are wider and much more frequent (min.: - 39.6°, max.: +45.9°, st.dev.: 20.3° with respect to t=0 μs), with rotations larger than 14.7° being observed in most of the sampled conformations. For VSD_I_, VSD_II_, and VSD_IV_, the channel simulated in a cholesterol-depleted raft-like membrane exhibits an intermediate behavior between the full raft-like and single-component membranes. VSD_III_ deviates from this trend, by showing overlapping behavior between the cholesterol-depleted and single-component membrane simulations (**Fig. 3d**).

### Tilt of VSD_IV_

A thinner membrane can cause the transmembrane (TM) helices to tilt, to align their vertical component with the thinner double layer. Across 10 of the 24 TM helices, the median tilt is different in the raft-like vs. single-component comparison. In particular, 7 of the TM helices exhibit a greater tilt in the single-component membrane, especially in DIII and DIV (**Fig. 3ef)**. Due to the preference for tilting in these domains and the lack of stabilizing effects from cholesterol and gangliosides at the region between PD and VSD_IV_, the latter domain shifts to establish tighter interactions with the DIII-IV linker (**Fig. 3f** and **S7**). Therefore, our simulations strongly indicate that the above-discussed modified Na_v_1.7 4-fold symmetry observed in the single-component membrane is due to the rotation of VSD_IV_ around the PD, which can move either in the direction of VSD_I_ or VSD_III_. Given the tight association of VSD_IV_ with the binding site of the IFM motif and its importance during the last stage of activation, this could have a great impact on the channel’s dynamics. Additionally, in the comparison between the cholesterol-depleted vs. full lipid raft-like membrane 4 TM helices display different tilting, with only 3 being more tilted in the cholesterol-depleted environment (**Fig. 3e** and **S6**). This suggests that cholesterol and membrane thickness have a limited, direct impact on inducing TM helix tilting in our simulations.

### Effect on pore and fenestration size

In Na_v_ channels, the pore cavity just below the selectivity filter is known to be the binding site for local anesthetics and antiarrhythmics^29^. Along the conduction pathway, at the cytosolic site of the inactivation gate, there is also the BIG (Below Inactivation Gate) site, where drugs like carbamazepine bind^30^. Side openings of the channel, referred to as “fenestrations”, are not only an access door for many molecules to the channel pore^29,31^, they are interaction hotspots for pyrethroids, DDT, and phospholipids^32–34^. We measured the pore size along the conduction pathway and through the fenestration, to test whether membrane composition has an impact on conduction or druggability of the binding sites in the pore and fenestrations. We found that in the comparison lipid raft-like environment vs. single-component membrane, a simple composition of the leaflets does not alter pore size at the level of the selectivity filter and only slightly increases the inactivation gate, but it does have a relevant impact on the central cavity which is reduced by 29.3% in volume, from 373.93 ± 63.43 Å^3^ to 264.45 ± 74.19 Å^3^ (**Fig. 4ab**). Fenestration size is always reduced in the single-component membrane, especially in the case of the fenestration formed by DII and DIII which is 43% smaller in volume and 0.67 Å narrower at its choke point (see **Fig. 4cd** and **Table S1**). In the membrane resembling a cholesterol-depleted raft, the channel adopts intermediate values between those observed in the other two conditions for the radius of central cavity (**Fig 4b**), pore (310.74 ± 83.80 Å^3^) and fenestrations (**Fig. 4d** and **Table S1**) volume.

**Figure 4.**
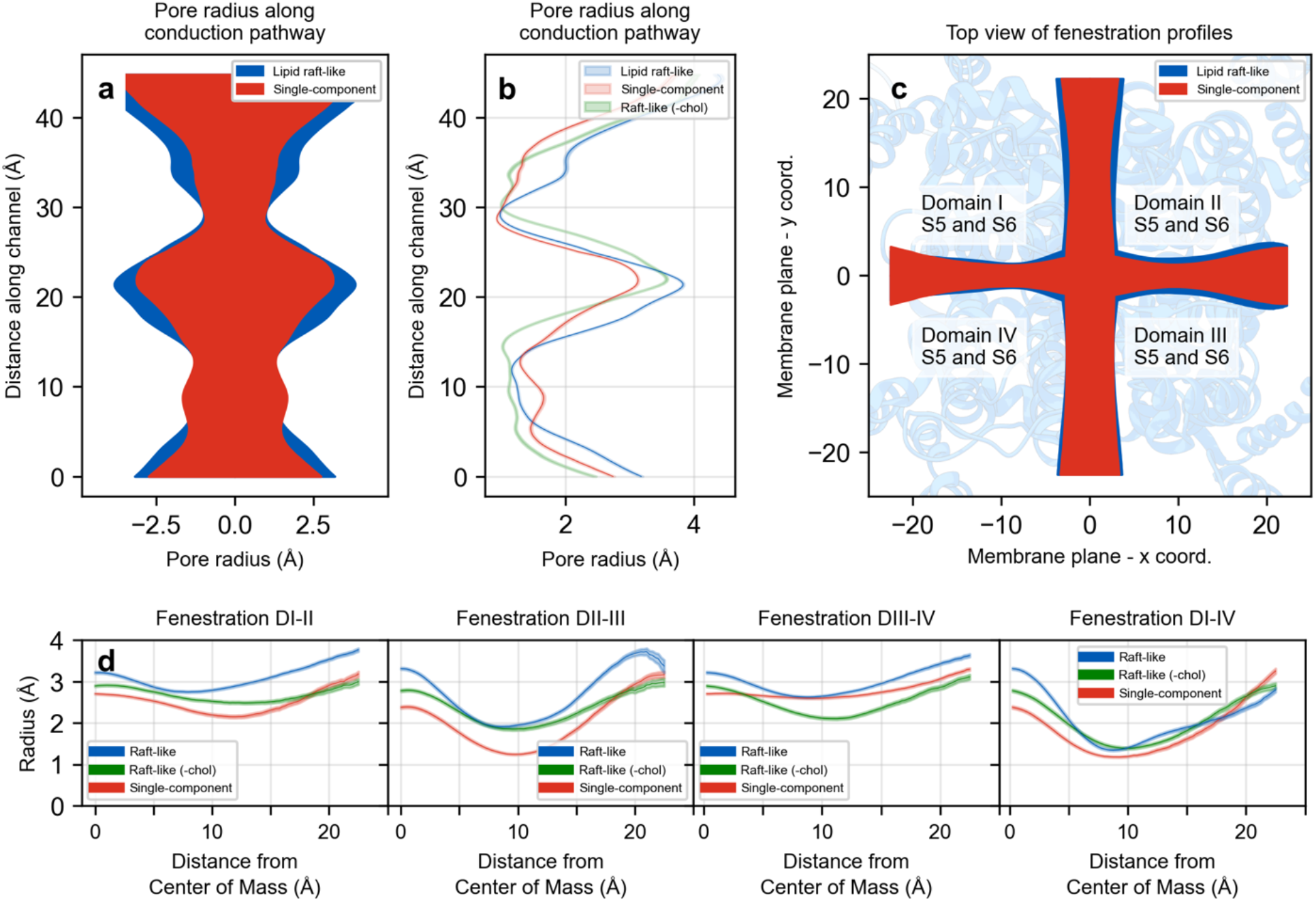
Changes in pore and fenestration size. Pore and fenestration size for the three membrane compositions (blue: lipid raft-like membrane; green: cholesterol-depleted raft-like membrane; red: single-component membrane). **a**. Side view of the pore profile along the conduction pathway. Green representation omitted for clarity (see panel b). **b**. Pore radius and bootstrapped 95%CI (n=1000 frames per simulation, representing 3 µs per membrane composition) along the conduction pathway. **c**. Extracellular view of fenestration profiles. Green representation omitted for clarity (see panel d). **d**. Fenestration radius and bootstrapped 95% CI along fenestration egress pathway.

### Effects on other binding sites

Traditionally, drug design targeting Na_v_1.7 involves two additional binding sites considered significant: the resting VSD_II_ and the activated VSD_IV_^35^. The former has been identified as a binding site for toxins like ProTx2 or m3-Huwentoxin-IV, while the latter can also interact with simpler organic compounds, such as aryl and acyl sulfonamides. Since ligand binding predominantly occurs extracellularly on the portion of the VSD facing away from the pore, we focused on monitoring significant changes in the shape on such binding domain by measuring the residue-to-residue distance between helices S2 and S3. In the VSD_II_, opposing changes were observed between the lipid raft-like and single-component conditions: in the lipid raft-like membrane, the extracellular ends of S2 and S3 move apart, while they tend to approach each other in the single-component membrane simulations (**Fig. S8**). In the membrane resembling a cholesterol-depleted raft, there is no consistent trend observed across the three replicas. However, in two of them, these ends slightly diverge. In the VSD_IV_, changes were more subtle, with only a trend of helices S2 and S3 slightly converging observed in the lipid raft-like membrane (**Fig. S9**).

Summarizing, our simulations showed an increase in the flexibility and tilting of VSD_IV_ in a single-component membrane and also an increase in the occupancy of cholesterol with VSD_III_. VSD_IV_ has been implicated in the fast inactivation process, while some studies have shown that the VSD_III_ can affect both the activation and fast inactivation processes^36^.

### Cholesterol depletion alters Na_v_1.7 activation and fast inactivation properties *in vitro*

Our simulation results suggest a possible role for membrane composition in regulating the activation and fast-inactivation processes of Na_v_1.7. We also observe *in silico* a high occupancy of cholesterol in regions important for gating. Combined with the known effects of cholesterol on gating of ion channels^37,11,9^, we hypothesized that cholesterol in the complex membrane likely plays a pivotal role in our observed results. To validate these hypotheses *in vitro*, activation and fast inactivation properties were measured using electrophysiology. To this end, whole-cell patch clamp of HEK293t cells transfected with human Na_v_1.7 wild type (hWT) plasmids was performed in control conditions (hWT ctrl) and under cholesterol depletion conditions using methyl-ß-cyclodextrin treatment (hWT MßCD). Cholesterol depletion was achieved by incubating the cells for 1h with the compound prior to recordings. MßCD is commonly used for the manipulation of cholesterol in various expression systems^38–40^. This compound has also been used to study voltage-gated sodium channels (e.g. Na_v_1.8), and in particular Na_v_1.4 in the same expression system used in this study^37,11^.

### Cholesterol depletion increases current densities and hyperpolarizes voltage-dependence of activation and steady-state fast inactivation

Under both control (without cholesterol depletion, defined as hWT ctrl) and experimental condition (with cholesterol depletion using MßCD, defined as hWT MßCD), robust inward currents were observed with quick inactivation kinetics (**Fig. 5a**). Patched cells showed a similar range of membrane capacitance values (C_slow_), which give an indirect measure of cell size (**Fig. S10**). To account for effects of cell size on the magnitude of inward currents, the currents were normalized to C_slow_ to obtain the current densities. The current densities of hWT MßCD are higher than the current densities of hWT ctrl (**Fig. 5b**). The maximal current densities of hWT MßCD (645.8 ± 186 pA/pF, n=15) were significantly higher than hWT ctrl (206.3 ± 89 pA/pF, n=14), with the average value 3.1 times higher than hWT ctrl (**Table 2, Fig. 5c; p < 0.05**). Persistent currents were measured as the mean current values between 34 ms and 39.6 ms of each test pulse and normalized to the peak currents. The maximal value for each cell was used for further analyses and depicted as a percentage of the peak currents. There was a small, but statistically significant decrease in the mean normalized persistent current values of hWT MßCD (1.3 ± 1.0%, n=15) when compared to hWT ctrl (1.8 ± 0.6%, n=14) (**Table 2, Fig. 5d; p < 0.05**). Given the relatively low amplitudes of the non-normalized persistent currents in hWT ctrl and hWT MßCD (**Fig. S11ab**), we checked if the values are higher than the noise in our system, which we assessed via the leak currents. The average persistent currents amplitudes were lower than the average leak current amplitudes in hWT ctrl and hWT MßCD across all voltage ranges (**Fig. S11cd**). Thus, while statistical significance is reached, the measure may be affected by the leak currents. Noticeable differences were measured in the voltage-dependence of activation and steady-state fast inactivation. hWT MßCD induced hyperpolarizing shifts of 7.2 mV and 7.8 mV in the V1/2 of activation (**Fig. 5de**) and steady-state fast inactivation compared to hWT ctrl, respectively (**Fig. 5fg**). However, no differences were observed in the slope of the curves (**Table 2**).

**Table 2:**
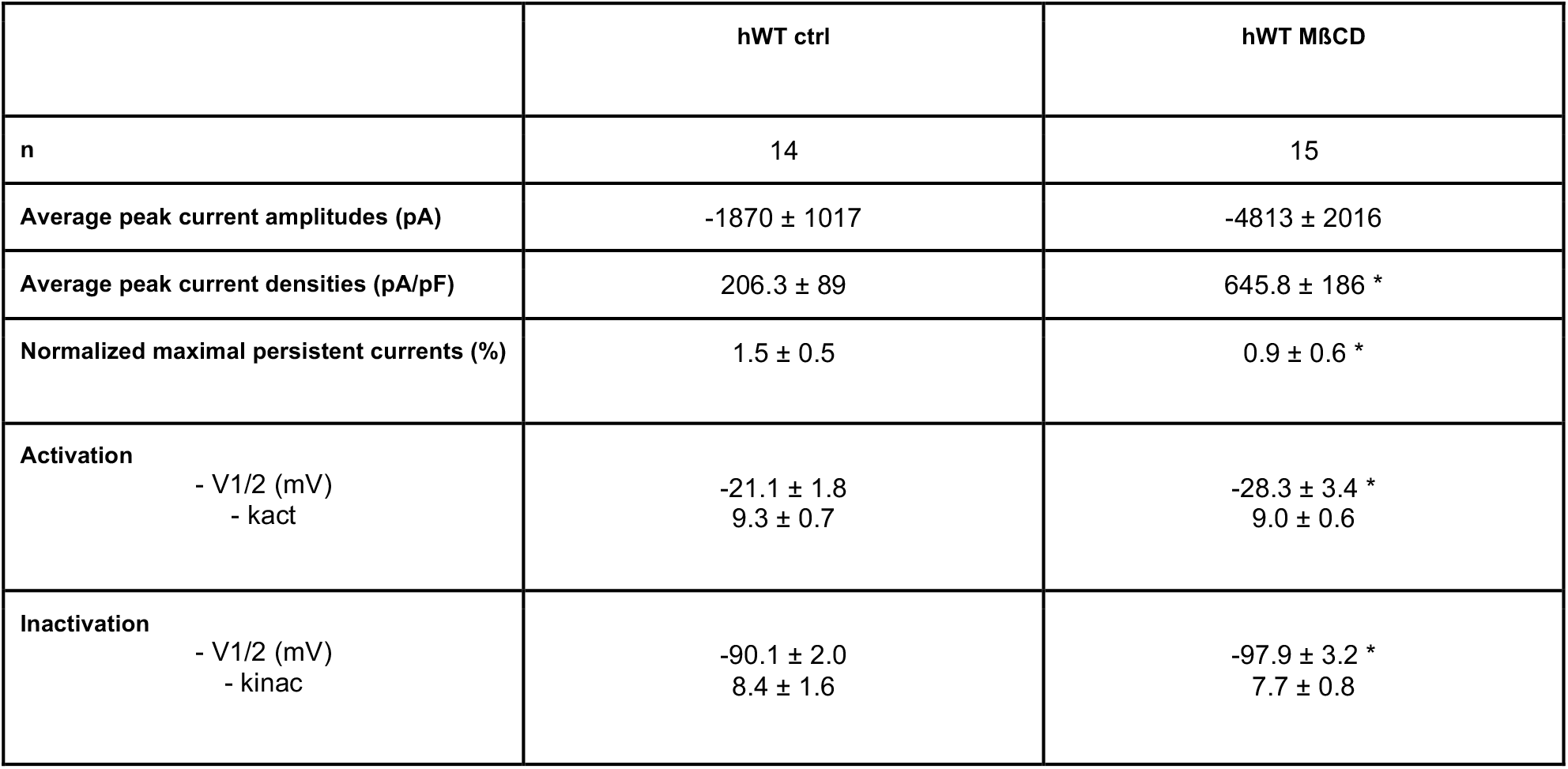
Current density, persistent currents, voltage dependence of activation and steady-state fast inactivation of human Na_v_1.7 wildtype under control (hWT ctrl) and cholesterol depletion (hWT MßCD) conditions. Values are represented as mean ± 95% confidence interval of the mean. Values of hWT MßCD that are significantly different from hWT ctrl are represented by a * in the hWT MßCD column for the parameters which underwent a statistical significance test and had a p value is less than 0.05.

**Figure 5:**
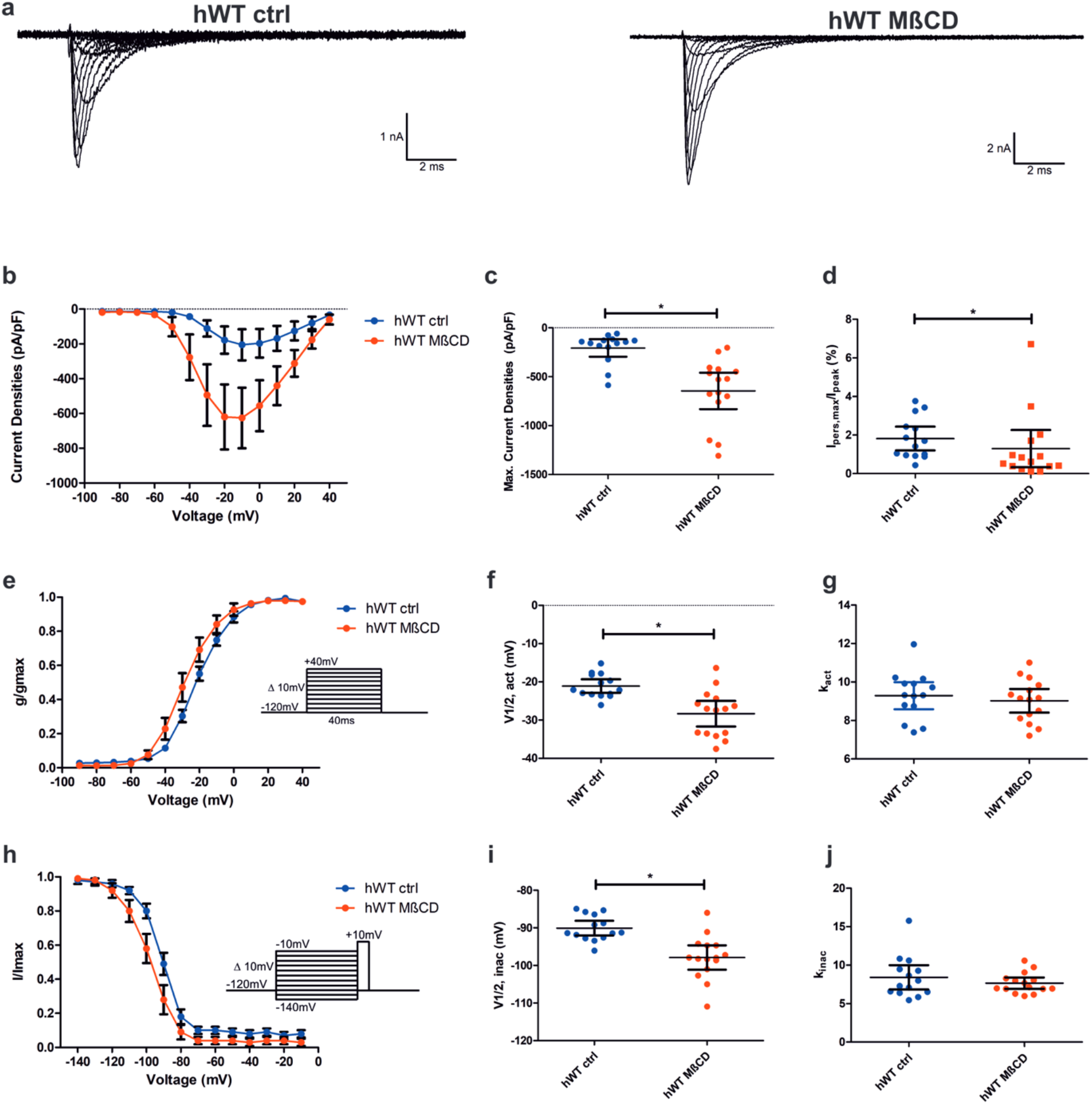
Current density and gating properties of human Na_v_1.7 wild type (hWT) under control and cholesterol depletion conditions. **a**. Representative traces of hWT control (hWT ctrl, left) and cholesterol depletion (hWT MßCD, right) conditions. Both conditions had robust inward currents. **b**. Average current density vs. voltage for hWT ctrl and hWT MßCD. Error bars represent the 95% confidence interval. **c**. Dot plot of peak current densities for hWT ctrl and hWT MßCD. Error bars represent the 95% confidence interval. * indicates a p-value < 0.05. **d**. Maximal persistent currents as a percentage of the peak transient currents (Ipers,max/Ipeak) vs. voltage (mV). Error bars represent the 95% confidence interval. * indicates a p-value < 0.05. **e**. Normalized conductance-voltage relationships of hWT ctrl and hWT MßCD. g is the conductance at a specific voltage and gmax is the maximum conductance. The voltage-dependence of the activation protocol is indicated in the inset. **f**. Dot plot of the V-half of activation for hWT ctrl and hWT MßCD. Error bars represent the 95% confidence interval. * indicates a p-value < 0.05. **g**. Dot plot of the slope factor kact of the G-V curve for hWT ctrl and hWT MßCD. Error bars represent the 95% confidence interval. **h**. Normalized current-voltage relationships of hWT ctrl and hWT MßCD. I is the inward current at a specific voltage and Imax is the maximal inward current. The voltage-dependence of steady state fast inactivation protocol is indicated in the inset. **i**. Dot plot of the V-half of steady state fast inactivation for hWT ctrl and hWT MßCD. Error bars represent the 95% confidence interval of the mean. * indicates a p-value < 0.05. **j**. Dot plot of the slope factor kinac of the I-V curve for hWT ctrl and hWT MßCD. Error bars represent the 95% confidence interval.

### Cholesterol depletion alters the time to peak and rate of onset of fast inactivation but not recovery from fast inactivation

In addition to voltage dependencies of activation and fast inactivation, the kinetics of these processes were also measured. The traces obtained with the activation protocol (**Fig. 5e** inset) gives us information on the onset kinetics of activation and fast inactivation. The time to peak is measured as the time it takes from the start of the stimulus pulse (**Fig. 6a** blue arrow) to reach the peak current (**Fig. 6a** green arrow). The time to peak was more rapid in hWT MßCD when compared to hWT ctrl for pulses between −50 mV and +40 mV (**Fig. 6b**; p < 0.05). The rate of onset of fast inactivation was measured by fitting the inactivating phase of the IV traces to a single exponential fit (**Fig. 6c** in red) and obtaining the time constant(s) of the fit(s). Cholesterol depletion decreased the time constants for fast inactivation leading to a quicker rate of onset of fast inactivation between −40 mV and +40 mV (**Fig. 6d**; p < 0.05).

**Figure 6:**
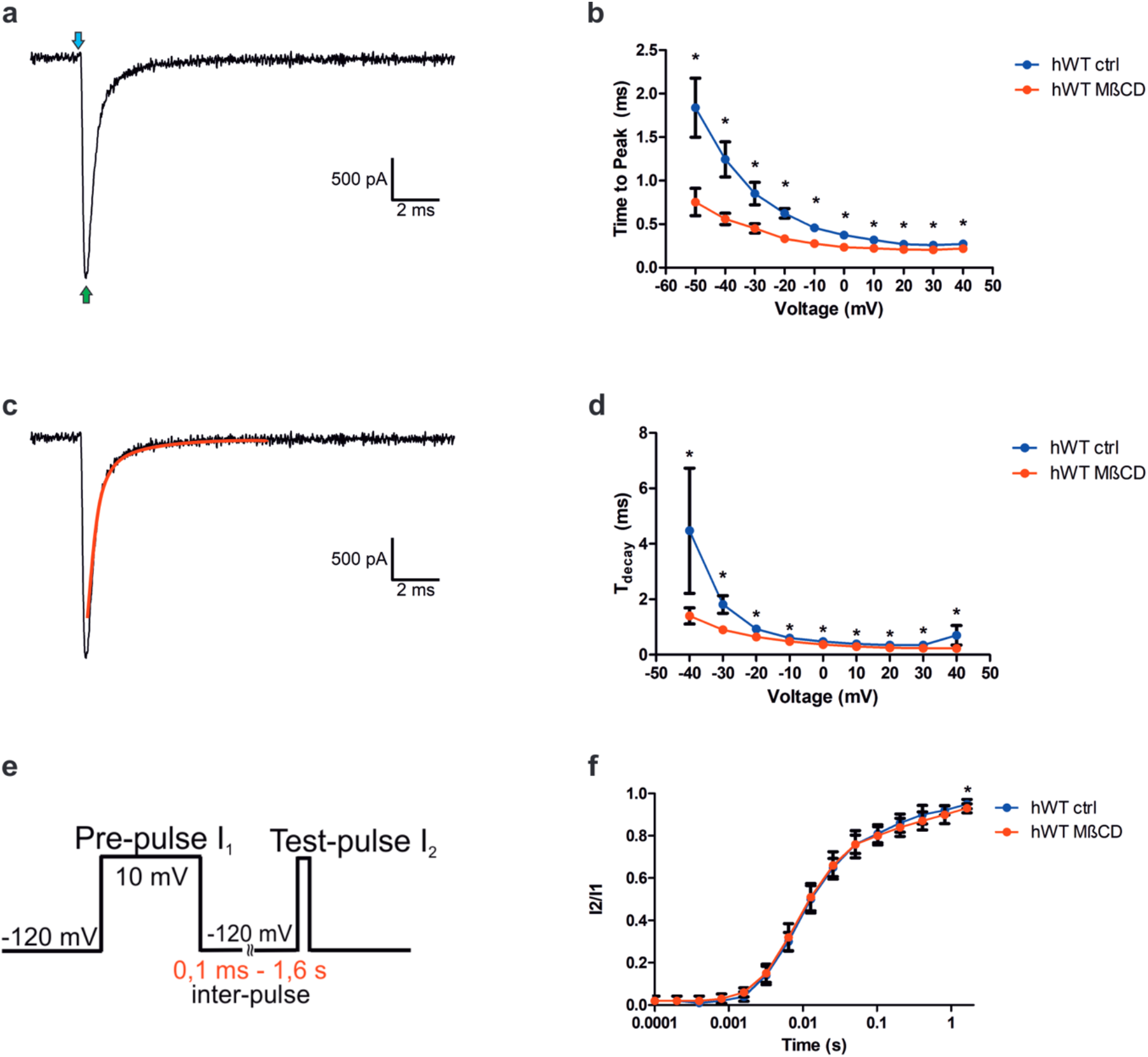
Kinetics of activation and fast inactivation. **a**. Calculation of time to peak. The time to peak is calculated as the time taken from the start of impulse (blue arrow) until peak current is reached (green arrow). **b**. time to peak vs voltage curves for hWT ctrl and hWT MßCD. * indicates a p-value < 0.05. Error bars represent the 95% confidence interval. **c**. Calculation of the time constant for decay of fast inactivation. The inactivating portions of the traces from the activation protocol (highlighted in red) are fit with a single exponential function. The time constant of the fit is used for analysis **d**. Time constant vs. voltage curves for hWT ctrl and hWT MßCD. * indicates a p-value < 0.05. Error bars represent the 95% confidence interval. **e**. Protocol to study the recovery kinetics from fast inactivation. **f**. Time course of recovery from fast inactivation. Peak currents from the test pulse (I2) were normalized to the peak currents during the pre-pulse (I1) and plotted against the duration of the interpulse. Error bars represent the 95% confidence interval of the mean.

The time course of recovery from fast inactivation was measured using the protocol shown in Figure 6e. The peak currents obtained in test pulse I2 were normalized against the peak currents obtained in pre pulse I1 and plotted against the duration of the interpulse. This yields a measure of how many channels recovered from fast inactivation (and hence are ready for subsequent activation in the test pulse) as a function of the time given for the channels to recover. Cholesterol depletion did not alter the recovery kinetics of fast inactivation except at an interpulse duration of 1.6s (**Fig 6e,f**).

## Limitations of the models

### In silico

All-atom MD simulations might appear as a preferable choice with respect to coarse-grained ones. However, the inherent slow dynamics of lipids, coupled with their intricate interplay within complex membranes and subtle changes in protein structure, pose challenges in obtaining adequate sampling. To investigate the influence of membrane composition on the dynamics of the Na_v_1.7 channel *in silico*, we therefore employed coarse-grained simulations with the Martini 2.2 force field. The latter is renowned for accurately simulating membrane proteins and their interactions with lipids^41,42^, despite the well known limitations of the forcefield^43^. In addition, the simulation protocols possess an inherent bias towards the initial protein configuration, due to the presence of an elastic network connecting the beads representing the protein’s backbone in the simulations. This means that large conformational changes cannot be observed in unbiased simulations. To avoid introducing artificial differences, we employed a consistent elastic network to preserve channel shape in all simulations. This ensured that observed variations arose from membrane composition changes rather than a different protein parametrization.

Another point to note is the deliberate exclusion of the N- and C-terminal regions along with the DI-DII and DII-DIII linkers from the Na_v_1.7 structure. This was done since these regions are exclusively intracellular and our focus was solely on the transmembrane regions of the channel. Moreover, the Na_v_1.7 structure used did not have good density maps in these regions due to their inherent flexibility^44^. Modeling intracellular linkers presents substantial challenges, whether through homology modeling or utilizing AlphaFold (alphafold.ebi.ac.uk/entryQ62205).

Lastly, it is important to note that we employed Martini 2.2 instead of the latest version, Martini 3, as the coarse-grained model in this study. Although Martini 3 has demonstrated notable improvements, this new version has not undergone comprehensive benchmarking for investigations involving protein-lipid interactions yet. In addition, Martini 3 protein and lipid models are still under development, with some bonded terms still not fully updated and a few important biomolecule models missing^45^. Recently, models for cholesterol^45^ and GM3^46^ were introduced, but the GM1 lipid model is still pending release. Given these considerations, we opted to conduct the study using Martini 2.2.

### In vitro

In the *in-vitro* approach, we used HEK cells to study the interplay between membrane composition and Na_v_ functionality. The plasma membranes of these cells show higher rigidity at induced high cholesterol concentration^18,47^, and, in standard conditions, different levels of sphingomyelin and phospholipids compared to other widely studied cell types^48^. Although a direct comparison of HEK cell membrane composition with sensory neurons is not reported, a recent publication investigates rat hippocampal neurons and suggests, similar to the results reported for HEK cells, that cholesterol enhances membrane rigidity, which may affect peptide insertion into the membrane^47^.

## Discussion

Cholesterol can have varying effects on different ion channels. For example, K_v_1.3 channels show inhibition of function upon increase in cholesterol concentrations, without any significant changes during depletion of cholesterol^49,50^. In contrast, voltage-gated calcium channels showed an increase of current densities and a hyperpolarizing shift in the voltage dependence of activation^51^, upon cholesterol depletion. Effects on membrane protein functioning by altered membrane properties have also been shown for mechanosensitive piezo channels^52^.

In the present study we show that cholesterol affects shape and channel gating of the pain-related sodium channel Na_v_1.7 by coarse-grain *in silico* modeling and *in-vitro* patch-clamp analysis of heterologously expressed channels.

The following were the major results from the study: **(1)** single component POPC membranes had reduced thickness and increased area per lipid compared to a full lipid raft-like membrane, consistently to what observed in other *in-silico* and *in-vitro* studies^19–21^; **(2)** *in silico* modeling of a cholesterol-rich lipid raft-like membrane increased the rigidity of the channel compared with the single-component POPC membrane, as emerging from decreased movements of the channel’s VSD_Iv_ and alteration of various of its geometric properties (i.e., pore and fenestration size); **(3)** *in vitro* cholesterol depletion increases current densities of Na_v_1.7 currents; **(4)** hyperpolarizes the voltage-dependence of activation and steady-state fast inactivation, and accelerates the time to peak and fast inactivation onset kinetics; **(5)** cholesterol in complex membranes is observed to frequently interact with the Na_v_1.7 protein close to the S4-S5 linkers *in silico*.

The effect of cholesterol on Na_v_ geometry and gating observed *in silico* and *in vitro*, respectively, could be attributed to indirect effects on the bilayer properties, or direct interactions with the channel protein^6^.

Indirect effects of the presence of cholesterol are usually mediated by an increase in the stiffness of the bilayer^6,53^, a reduction in its fluidity^18,47^, and the resulting denser arrangement of the lipids^54^. Our first finding is in agreement with such experimental observation (**Finding 1**). Cholesterol-driven membrane’s stiffness can indirectly impact on the channel dynamics: Na_v_ channel activation indeed involves a pronounced movement of the S4 helix within the voltage-sensor and, via displacement of the S4-S5 linker, it induces dilation of the activation gate in the pore allowing ion permeation^55–57^. Similarly, fast inactivation is initiated by displacement of the DIII-DIV linker containing the inactivation particle, which finally binds to the side of the channel pore, thereby closing the permeation pathway. Thus, both, activation and fast inactivation, are based on larger movements within the channel protein, and a less rigid membrane surrounding may result in speeding of these processes. Notably, larger rototranslational movements of the VSD_IV_ are observed *in silico* when cholesterol is absent, and these allow a tighter association to the inserted DIII-DIV linker (**Finding 2**). Consistently, in our patch-clamp recording in Na_v_1.7 expressing cholesterol depleted HEK cells, we observed a speeding of time to peak and fast inactivation onset kinetics. Also steady-state fast inactivation and channel activation occurred at more negative potentials, indicating that less energy is needed for these gating processes when cholesterol is depleted. The observed increase in current density may result from quicker, and thus more coordinated channel opening. Alternatively, the reduced rigidity of the membrane may result in a higher rate of channel insertion, resulting from facilitated trafficking. Thus, our findings suggest that the observed effects on channel gating are probably due to indirect action of cholesterol depletion on the properties of the membrane (**Finding 3 and 4)**.

Conversely, the clustering of cholesterol molecules around the S4-S5 linkers observed *in silico* suggests that cholesterol can also directly have an impact in the channel’s activation by restricting the displacement of the S4-S5 linker (**Finding 5**). The latter movement is necessary for the activation of the VSDs and the opening of the activation gate in the pore allowing ion permeation.

Comparing our findings with what was previously observed for other Na_v_s, cholesterol depletion appears to affect these channels in a subtype specific manner. On one hand, unlike our study, cholesterol depletion of Na_v_1.4 in HEK293 cells did not increase current densities or change activation properties^37^. On the other hand, in agreement with our studies, Na_v_1.9 showed an increased activity upon cholesterol depletion that was attributed to direct effects of cholesterol upon binding to any of the 17 motifs identified in the study^9^. Cholesterol depletion in Na_v_1.8 also possibly reduced the channel’s surface expression and reduced the number of neurons conducting depolarizations^11^.

Changes in membrane properties might be a more general mechanism by which cholesterol depletion alters the gating of Na_v_s. Other indirect effects such as altered interactions with signaling molecules, e.g. kinases, and direct binding of cholesterol to regions in the channel, may underlie subtype specific differences of cholesterol depletion and present a promising avenue for further research.

In a broader view, our findings highlight the importance of evaluating the effects of membranes when employing structure-based drug discovery for ion channels, as they can impact the size and shape of relevant binding sites. Specifically for Na_v_1.7, our study opens up new paths for rational drug design of novel analgesics that take into account cholesterol modulation and the environmental features of the channel’s localization.

## Material and Methods

### Pairwise sequence alignment

Mouse Na_v_1.9 (uniprot ID: Q9R053) and human Na_v_1.7 (uniprot ID: Q15858-1) sequences were aligned using the online tool EMBOSS Needle. The latter implements a EBLOSUM62 matrix with a gap penalty of 10.0, and an “extend” penalty of 0.5 (https://www.ebi.ac.uk/Tools/psa/emboss_needle/).

### Molecular Dynamics

All simulations were performed using Gromacs 2019.4^58,59^ and the Martini 2.2 forcefield with ElNeDyn22 for protein, water and ions, and, and lipids 2.0^60-63^ for lipids. The structure of the channel was downloaded from the PDB database (PDB ID: 6J8G)^44^; loops were repaired using Modeller 9.10^64,65^, aligned vertically to a horizontal membrane bilayer using PPM webserver 2.0^66^. The topology of the protein was created using Martinize^63^. The default settings for the elastic network were used, with a final number of 4385 distance restraints, with minimum and maximum distance of 0.33 nm and 0.9 nm, respectively. The elastic bond force constant for all the bonds was set to 500 kJ/mol/nm^2^. The protein was embedded in the membrane using Insane (https://github.com/Tsjerk/Insane). The raft-like membrane was taken from the MERMAID webserver representation of a human myelin membrane^67^ and it is formed by 12% phosphatidylcholine (POPC), 18% phosphatidylethanolamine (POPE), 30% Cholesterol (CHOL), 10% phosphatidylserine (POPS), 1% phosphatidylinositol (POPI), 8% sphingomyelin (POSM) and 21% monosialotetrahexosylganglioside (DPG1). In the raft-like (-chol) membrane, the cholesterol is replaced by POPC, while in the single-component membrane, the bilayer is composed of 100% POPC. The salt (NaCl) concentration was set to 0.15 M. The box size containing the system was adjusted with a short 2 ns NPT simulation, then the system was equilibrated with 200 ns of simulation in an NVT ensemble, followed by 200 ns in an NPT ensemble. During equilibration, 1000 Kj/mol restraints were applied to the backbone of the protein. For each condition, three 4-µs replicas were produced.

### Domain rotation analysis

The intervals defining the 24 transmembrane helices were obtained from Uniprot (gene: SCN9a_human, ID: Q15858) and adapted to match isoform 3 of the channel. The center of mass of each VSD was calculated using the masses of the helices S1 to S4. To calculate the rotation of the VSD around its vertical axis, a vector connecting the S1+S2 and S3+S4 centers of mass was defined for each domain. An analysis of the distributions of these angles over time is detailed in **Fig. S12** and **S13**.

### Pore and fenestration size

The program HOLE 2 was used to measure conduction pore and fenestration size of the coarse-grained channel during the last microsecond of production run (1 frame/ns, 1000 frames per simulation) of all the replicas^68^. The protein was roto-translated to fit the starting configuration using GROMACS 2019.4. The starting point for the sampling was set to the center of mass of the backbone. The conduction pore was measured along the z coordinate of the simulation box; whereas the fenestration direction was defined by the two vectors connecting residue 952 to 1634 and residue 249 to 1441. The limits of the central cavity were defined on the first frame from the z coordinate of the selectivity filter (5.8 Å above the center of mass of the protein) and Tyrosine 1755 (Na_v_1.7 isoform 1 numbering) (12.4 Å below the center of mass of the protein). The calculations for each fenestration were truncated at a distance of 22.5 Å from the center of mass of the protein. The null values rarely obtained in the calculations, represent an imperfection of the pathfinding algorithm that does not always succeed in finding a complete path from the begin to the end of the otherwise-open pore or fenestrations. Therefore, these values were excluded from the calculations, rather than being considered an interruption in the path. Van Der Waals radii were set to 2.35 Å for the standard beads, and 2.15 Å for the smallest ones. The sampled radii were extracted in Python 3.9.7, aligned, and averaged and plotted using the Numpy (v 1.21.2)^69^ and Matplotlib (v 3.4.3)^70^ libraries. Volumes were approximated as the sum of π x (sampled radii)^2^ x *h*, where h is the sampling interval corresponding to 0.25 Å. Volumes were calculated per each frame, and the average and standard deviation of 3000 frames per condition was reported. The chokepoint was measured as the minimum non-null radius in the fenestration. It was calculated per each frame and the average and standard deviation of 3000 frames per condition was reported. Bootstrapping of the radii was performed with the scipy (v 1.8.0) python library^71^ and “BCa” method and n=1000 and reported in the visualization in **Fig. 4**.

### Lipid bilayer thickness, area per lipid and cholesterol binding motif analysis

The position of the head of phosphatidyl lipids (beads named PO4) was extracted from the simulations via MDAnalysis (v 2.0.0) and split into upper and lower layer groups by a horizontal plane placed in correspondence of the average *z* positions of the beads. The subdivision in two layers was updated every frame. The distance between the two leaflets was sampled for all simulations with a sample rate of 1 sample/ns, for a total of 12,000 measurements per membrane-type. The distribution of the area per lipid was measured using Fatslim^72^ and by selecting all PO4, ROH, and AM1 beads as lipid heads. In this case too, the subdivision in two layers was updated every frame. All protein beads were treated as “interacting groups”. Emboss fuzzpro (www.bioinformatics.nl/cgi-bin/emboss/fuzzpro) was used to search for CRAC and CARC sequences in the SCN9A protein sequence isoform 1 reported in Uniprot (ID: Q15858-1 (www.uniprot.org/uniprotkb/Q15858/entry#Q15858-1), using as a query “[LV]X(1,5)YX(1,5)[RK]” and “[RK]X(1,5)YX(1,5)[LV]”.

### Analysis of transmembrane helices tilt

We utilized the centers of mass from the first and last four residues of every transmembrane helix to establish a vector that indicates the direction of the helix. TM helices regions were selected as discussed above. These vectors were then compared to the unit vector of the z-coordinate of the simulation box, and the resulting angle between them was recorded for each helix once every nanosecond. The average (last 1 µs, 1000 frames) TM helix tilt of the replicas were split into two groups representing the two conditions to be compared with an independent t-test. The significance threshold was set at p<0.05.

### Analysis of occupancy of membrane components

The analysis was performed with VMD (1.9.4a55) and the integrated VolMap tool^73^. For the calculation, a grid spacing of 1 Å was used and simulated particles were considered point-like. A total of 12,000 frames (1 frame/ns) per membrane composition was used for the analysis, representing 12 µs.

### Transfection of HEK293t Cells

HEK293t cells were cultured in were cultured in Dulbecco’s modified Eagle’s medium (DMEM/F12; Gibco-Life Technologies, Carlsbad, CA, USA), supplemented with 10% fetal bovine serum (FBS; Gibco-Life Technologies, Carlsbad, CA, USA) and incubated at 37°C and 5% CO_2._ Human Na_v_1.7 (hWT) plasmid was contained in a pCMV6Neo vector. 2-5h post seeding of the HEK293t cells, 1.25 µg of hWT plasmid and 0.25 µg of green fluorescence protein (pMax-GFP) plasmid were co-transfected into the cells using 3 µL of the jetPEI® transfection reagent (Polyplus-transfection S.A., Illkirch, France) and incubated overnight at 37°C and 5% CO_2_. Transfected cells were patched 24h post transfection.

### Manipulation of Cellular Cholesterol Content

Membrane cholesterol content was decreased by exposure to the cholesterol chelating agent methyl-ß-cyclodextrin (MßCD). 258.2 g of powdered MßCD (Sigma-aldrich, C4555) was dissolved in 1 mL of distilled water and vortexed to produce a 200 mM clear MßCD stock solution. The stock solution was mixed with DMEM/F12 medium to create a 5 mM MßCD medium. HEK293t cells that were used for cholesterol depleted conditions were incubated in the 5 mM MßCD medium for 1h at 37°C and 5% CO_2_ before patching.

### Whole-cell Voltage Clamp Experiments

Whole-cell voltage clamp recordings of transfected HEK293t cells were performed at room temperature using the HEKA EPC 10 USB amplifier (HEKA Electronics, Lambrecht, Germany).

The recording pipettes were pulled from borosilicate glass capillaries using a DMZ puller (Zeitz Instruments GmbH, Martinsried, Germany). The pipette resistances varied between 1-3MΩ. The external bath solution contained the following (in mM): 40 NaCl, 100 Choline-Cl, 3 KCl, 1 MgCl_2_, 1 CaCl_2_, 10 HEPES, 5 glucose and 10 sucrose. pH of the solution was adjusted to 7.4 using CsOH and osmolarity to about 298 mOsm. 1 in 1000 parts dimethylsulfoxide (DMSO) was added to the bath solution. The internal pipette solution contained the following (in mM): 10 NaCl, 140 CsF, 1 EGTA, 19 HEPES and 18 sucrose. pH of the solution was adjusted to 7.3 using CsOH and osmolarity to about 310 mOsm.

For all cells recorded and considered for further analyses, series resistance was compensated by at least 70% to ensure that voltage errors did not exceed 5 mV. Leak current was subtracted digitally using the P/4 procedure. Unless stated otherwise, the holding potential was maintained at −120 mV. No liquid junction potential corrections were performed. After whole-cell configuration is reached, the inward sodium currents generated by the hWT-expressing cells were allowed to stabilize for 3min during repeated −10 mV depolarization steps before starting the protocols.

The voltage-dependence of activation was assessed from the holding potential (Vhold) using 40ms test pulses with varying potentials from −90 mV to +40 mV in 10 mV increments, with a 5s inter-pulse interval (Figure 5e, inset). The inward currents (I) recorded at each voltage step (V) could then be used to calculate the conductance (G) by the equation: G = I/(V-Vrev), where Vrev is the reversal potential for sodium. The conductance-voltage curves could then be fit using a boltzmann equation to obtain the voltage-dependence characteristics of activation: G = Gmax/(1 + exp[Vm - V1/2)/k_act_]), where Gmax is the maximal sodium conductance, Vm is the membrane voltage, V1/2 is the potential of half maximal activation and k_act_ is the slope factor. The time course of fast inactivation, which estimates how rapidly channels fast inactivate, can be obtained for each test pulse by fitting the inactivating portion of the activation trace (**Fig. 6c**) to a single exponential function: Y = Y0 + A*exp(-KX), where Y0 is the steady-state current amplitude, A is the amplitude coefficient, K is the rate constant and X is the time. The time constant τ_decay_, which is the reciprocal of K, is plotted against the test-pulse voltages from −40 mV to +40 mV. The time to peak is measured for each test pulse by calculating the time taken between the start of the test pulse and the channels reaching Imax (**Fig. 6a**). The time to peak is plotted against the test-pulse voltages between −50 mV and +40 mV. Maximal current densities for each cell were calculated by normalizing the peak currents for each test pulse by the membrane capacitance of the cell. Current density values have the unit pA/pF.

The voltage-dependence of steady-state fast inactivation was measured using a two-step protocol. Firstly, a 500 ms pre-pulse from Vhold with potentials ranging from −140 mV to −10 mV in 10 mV increments was used to inactivate the channels. This was followed by a 40 ms test pulse to 0 mV to gauge the fraction of channels that are still fast inactivated (**Fig. 5h**, inset). The inward currents (I) measured during the test pulse were normalized to the maximal inward current (Imax) of the cell and plotted against the pre-pulse voltages (V) to obtain a normalized current-voltage curve. The curve was fit using the boltzmann equation: I/Imax = Imin + (Imax-Imin)/(1+exp[(V1/2 - V)]/k_inac_), where Imin is the minimal inward current of the cell, V1/2 is the potential of half-maximal channel fast inactivation, and k_inac_ is the slope factor.

Recovery from fast inactivation was also measured using a two-pulse protocol. Cells were first depolarized using a pre-pulse to +10 mV for 500 ms from Vhold. This was followed up by an inter-pulse to −120 mV with increasing durations in each sweep by a factor of 2 - from 0.1 ms to 1600 ms. The inter-pulse was followed up with another depolarizing test pulse to +10 mV for 40 ms to obtain the fraction of channels that could recover from fast inactivation (Figure 6e). The current amplitudes at the test pulse (Itest) were normalized to the current amplitude at the pre pulse (Ipre) and plotted against the inter-pulse durations to obtain the time course of recovery from fast inactivation.

### Whole-Cell Voltage Clamp Data Analysis and Statistics

Raw data of the recordings were processed using Fitmaster (HEKA Electronics, Lambrecht, Germany) and exported into Igor Pro (Wavemetrics, Portland, OR, USA) for extraction of features from the traces using in-house scripts. Curve fittings and generation of the graphs were done using Graphpad Prism 5 (Graphpad Software Inc., La Jolla, CA, USA). For statistical testing, the control and cholesterol depleted conditions were compared using either a Student’s t-test or a mann-whitney U test based on the normality of the data which was tested using the Shapiro-Wilk normality test. The data is always represented as the mean ± 95% confidence interval of the mean and the error bars in the graphs denote the 95% confidence interval of the mean.

