## Supplementary Information for "Depletion of membrane cholesterol modifies structure, dynamic and activation of Na_v_1.7"

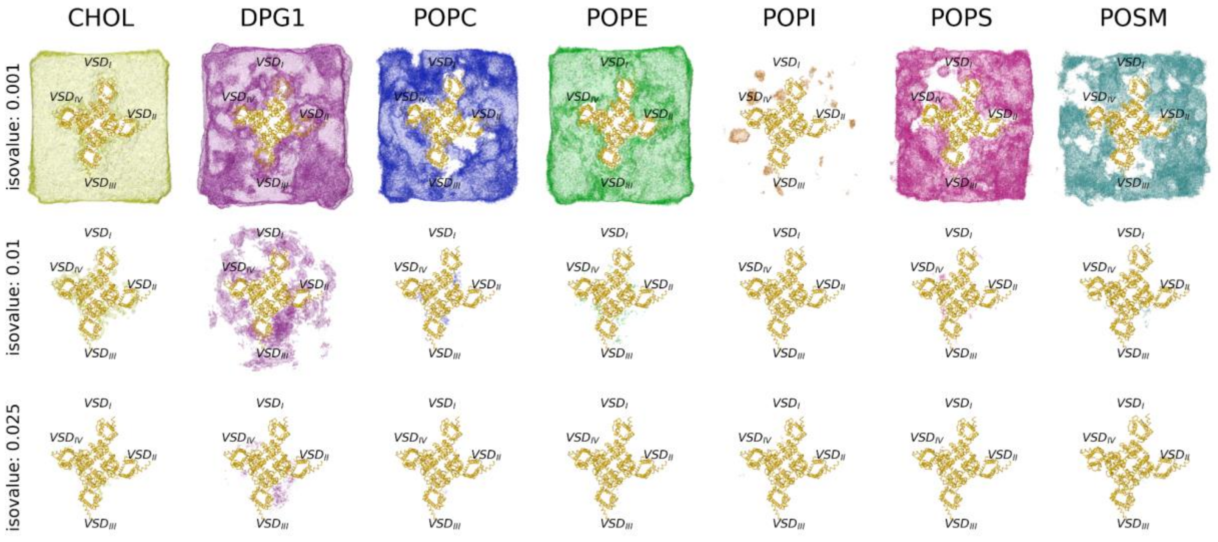


**Figure S1: Occupancy of lipids and cholesterol in the raft-like neuronal membrane**. The occupancy of each membrane component around the channel is represented as isosurfaces. The value of each 1-Å spaced grid point in the isosurface is set to either 0 or 1, depending on whether the corresponding grid cell contains one or more beads or not, at a given frame. The average of all the considered frames (12,000 frames, 1 frame/ns) provides the fractional occupancy of that point for the component. The surface is delineated by all points possessing a fractional occupancy value that matches the isovalue and encompasses every point with an occupancy value that is equal to or greater than this isovalue. In this figure, each row delineates the isovalues corresponding to thresholds of 0.1%, 2.5%, and 1%.

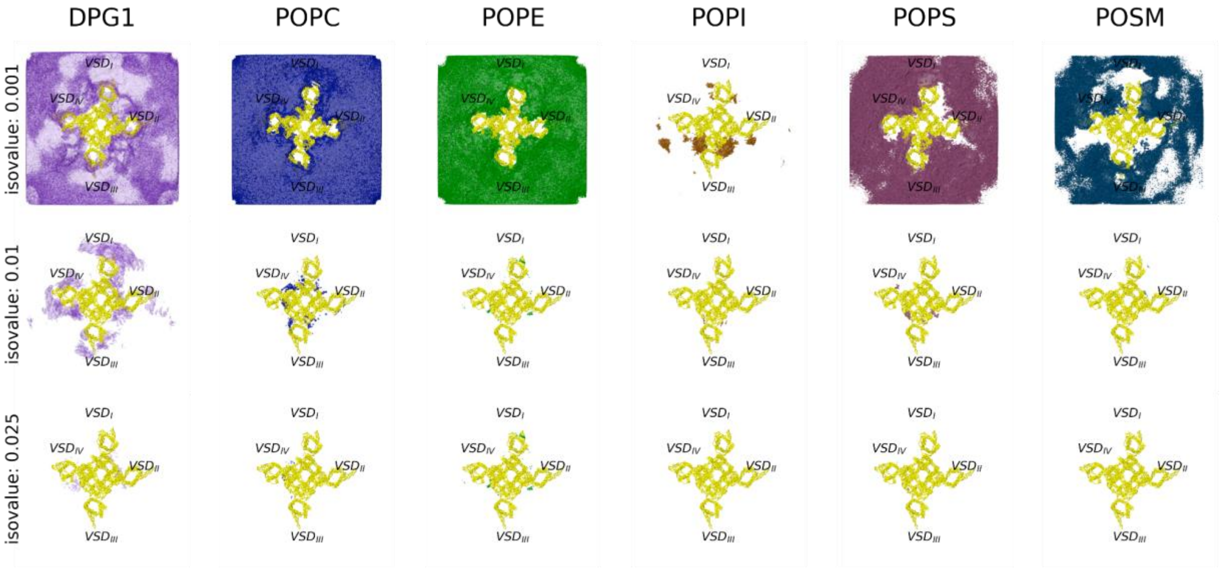


**Figure S2. Occupancy of lipids in the cholesterol-depleted raft-like neuronal membrane.** Lipids occupancy during the 12 µs (4 µs for each replica) of unbiased simulations in the lipid raft-like (-chol) neuronal membrane. The visualization technique employed here is consistent with the one delineated in Figure S1, with 12,000 frames analyzed, 1 frame/ns.

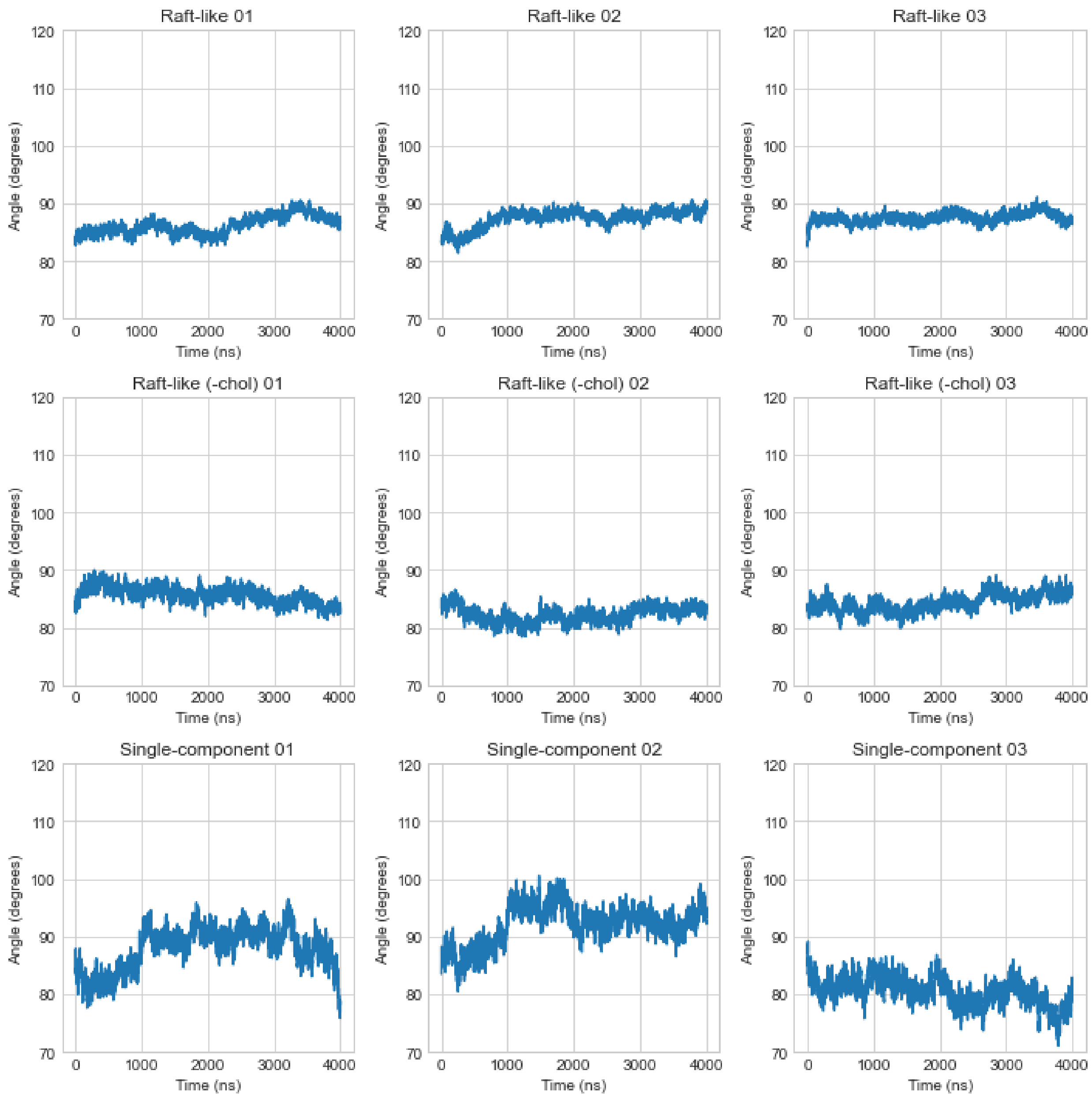


**Figure S3. Symmetry of the VSD around the pore.** Values of the angle between vectors connecting $VSD_{I}$ to $VSD_{III}$ and $VSD_{II}$ to $VSD_{IV}$ during 4 µs of unbiased simulation per replica.

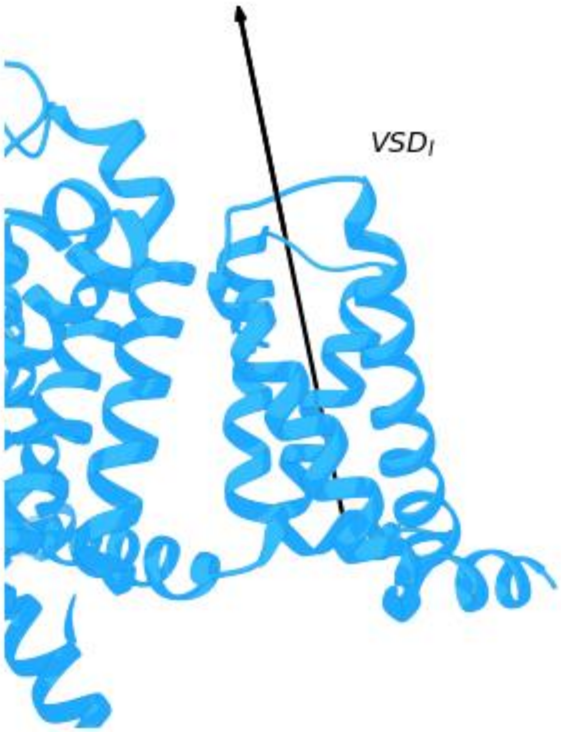


**Figure S4. Axis of rotation of VSDs.** The rotation around the axis (as defined in Figure 3c) defines the orientation of the VSD with respect to the pore domain.

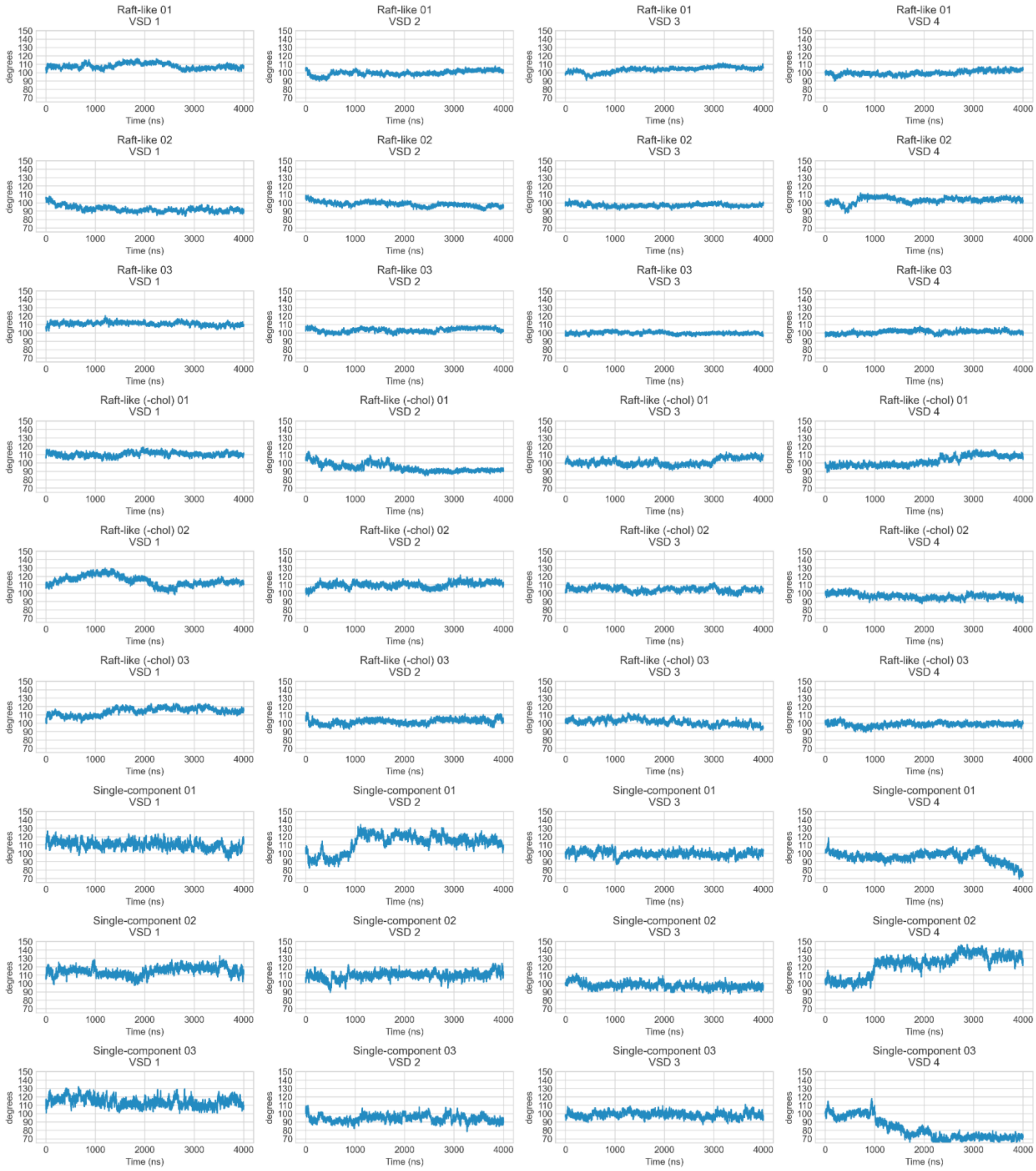


**Figure S5. Angle of rotation of VSDs**. VSD rotation around their intracellular-to-extracellular axis, during 4 µs unbiased simulation per each replica.

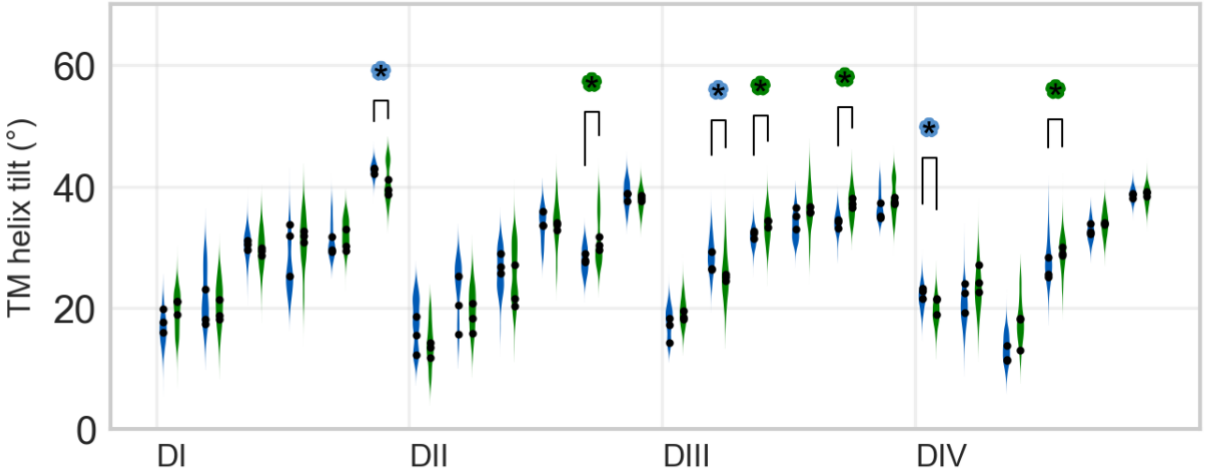


**Figure S6: TM tilt angle.** Distribution of tilt angle for the 24 TM helices in the lipid raft-like and cholesterol-depleted lipid raft-like membrane. The average of the three replicas per membrane composition is represented by black circles. A coloured stroked asterisk marks a significant difference in the tilt of an TM between the two conditions (see test in methods).

**Table S1: Fenestrations' size.** Analysis of the fenestration volume is reported for each membrane's composition, averaged over the last 3 µs across each replica (column 1 to 3). A comparison between the fenestrations' volumes on passing from raft-like vs. single-component is offered in column 4. All the analyses were conducted with *HOLE* software[1] (see methods for details).

| Fen. | Avg. and st.dev. of fenestration volume - Raft-like (Å3) | Avg. and st.dev. of fenestration volume - Raft-like (-chol) (Å3) | Avg. and st.dev. fenestration volume - Single-component (Å3) | % Reduction in volume in raft-like vs. single-component comparison |
|---|---|---|---|---|
| DI-II | 591 (100) | 443 (128) | 530 (71) | 10% |
| DII-III | 481 (129) | 412 (141) | 333 (100) | 31% |
| DIII-IV | 632 (133) | 506 (121) | 420 (119) | 33% |
| DI-IV | 368 (104) | 342 (89) | 264 (80) | 28% |
| Fen. | Fenestration choke point size and st.dev. - Raft-like (Å) | Fenestration choke point size and st.dev. - Raft-like (-chol)(Å) | Fenestration choke point size and st.dev. - Single-component (Å) | % Reduction in choke point size in raft-like vs. single-component comparison |
| DI-II | 2.08 (0.46) | 1.24 (0.57) | 1.98 (0.35) | 5% |
| DII-III | 1.21 (0.64) | 1.20 (0.77) | 0.71 (0.25) | 41% |
| DIII-IV | 2.18 (0.61) | 1.63 (0.65) | 1.27 (0.56) | 42% |
| DI-IV | 0.76 (0.17) | 0.68 (0.20) | 0.66 (0.17) | 12% |

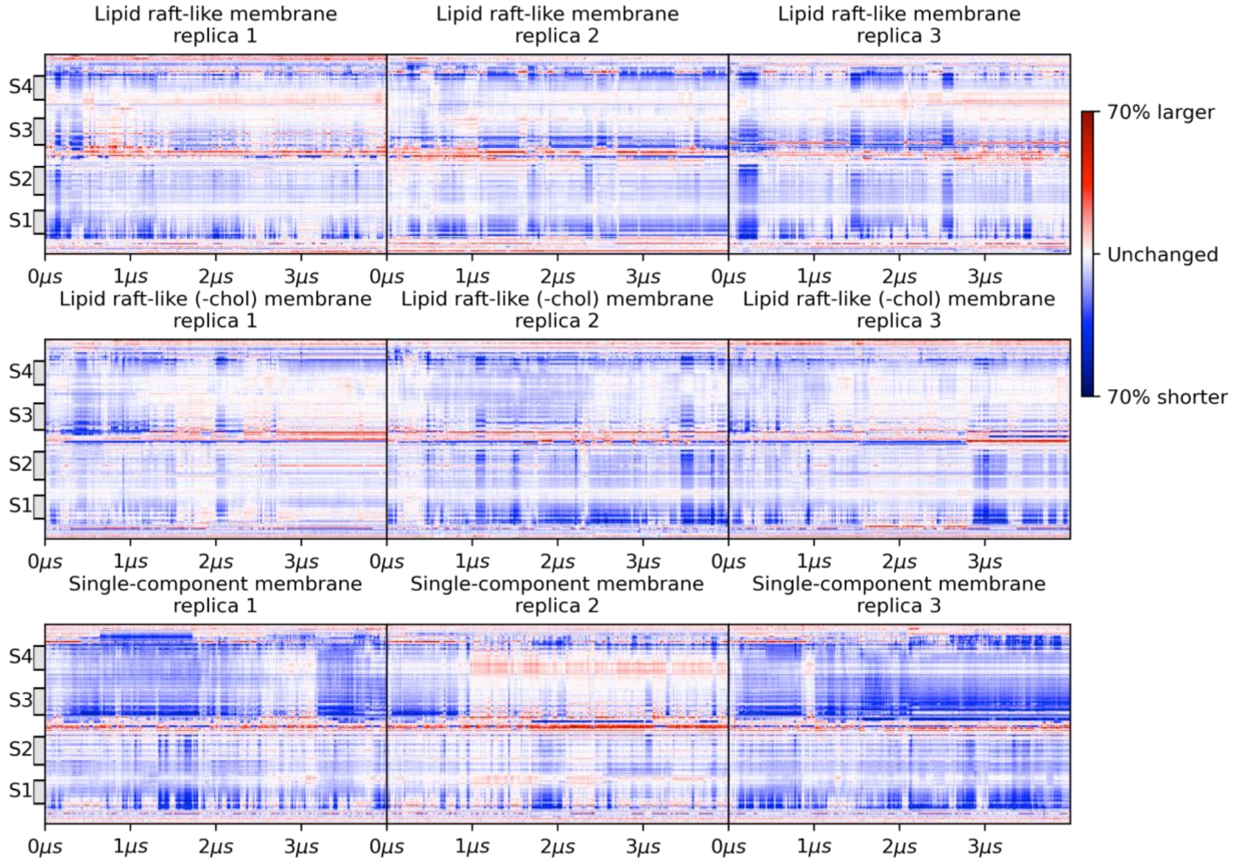


**Figure S7: VDS$_{IV}$ minimum distance to the DIII-DIV linker.** Variation in the distance between residues of the VSD$_{IV}$ (residues 1491-1630) to the ones of the DIII-DIV linker (residues 1456-1490) for each replica in each membrane composition. The channel conformation at t=0 μs is used as a reference.

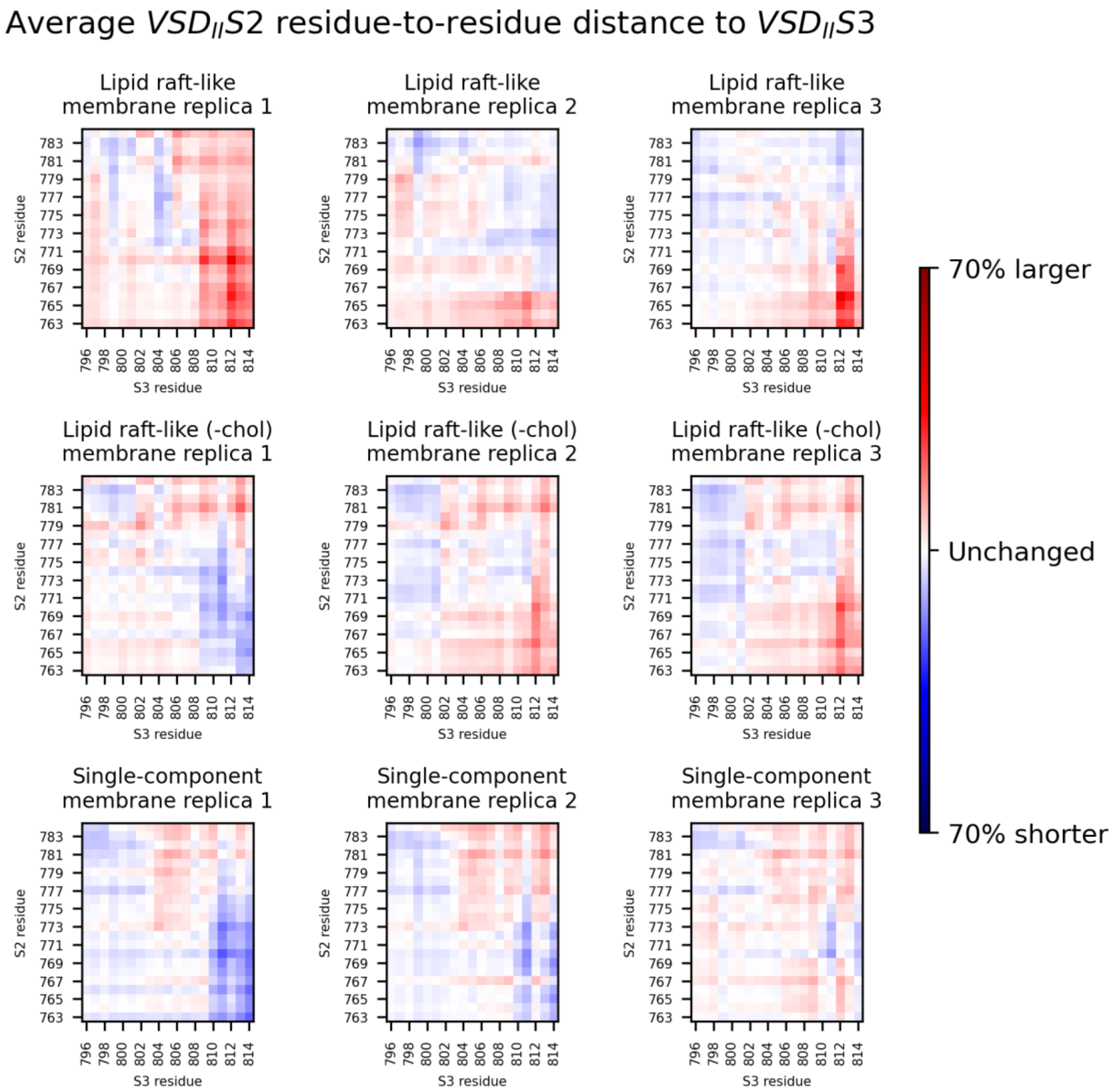


**Figure S8: Distance matrix between S2 and S3 in $VSD_{II}$.** Average residue-to-residue distance matrix between helices S2 and S3 in $VSD_{II}$ for each replica in each membrane composition. Distances were collected every 1 ns and normalized by the value at time t=0 µs.

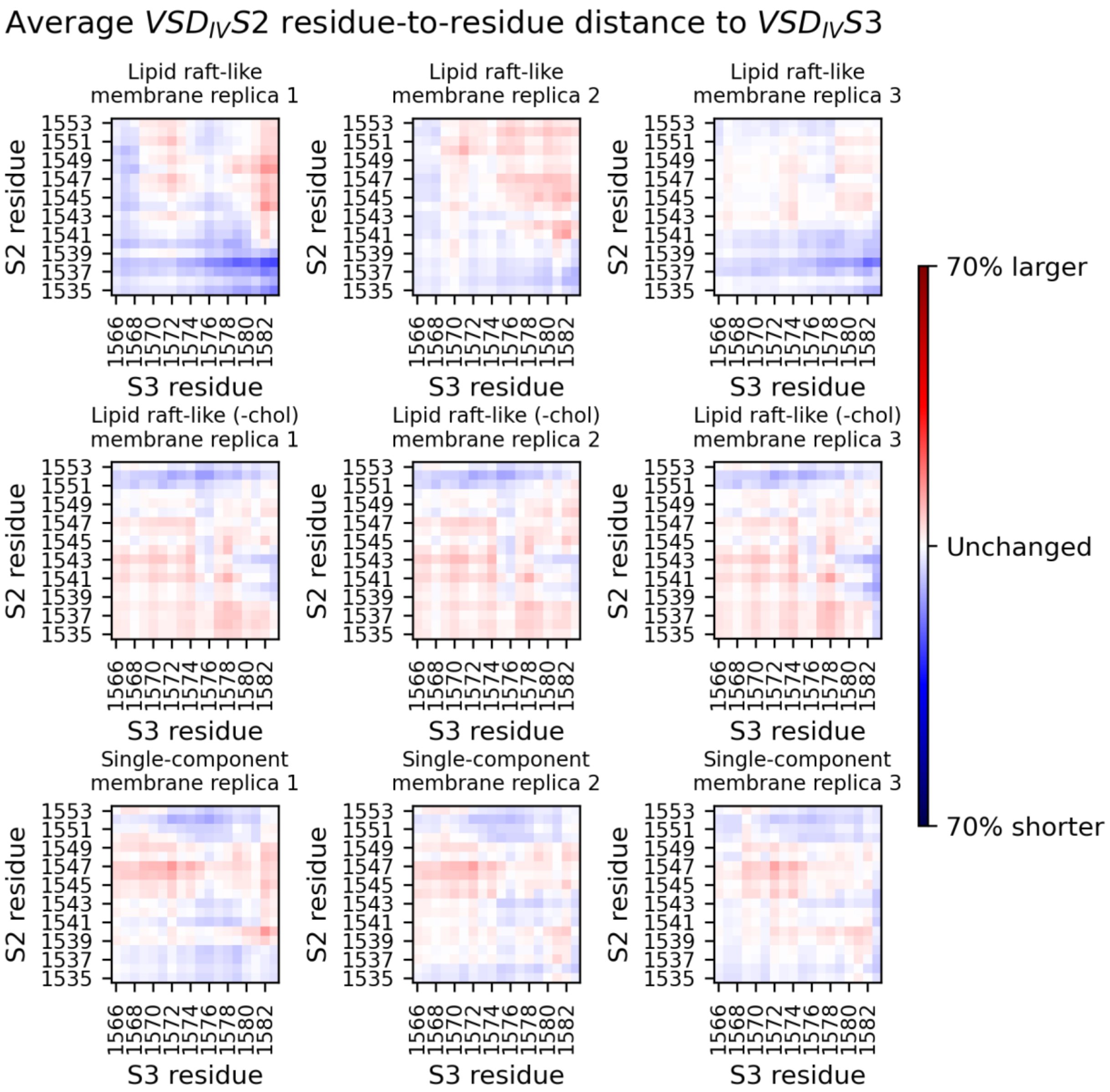


**Figure S9. Distance matrix between S2 and S3 in VSD$_{IV}$.** Average residue-to-residue distance matrix between helices S2 and S3 in VSD$_{IV}$ for each replica in each membrane composition. Distances were collected every 1 ns and normalized by the value at time t=0 µs.

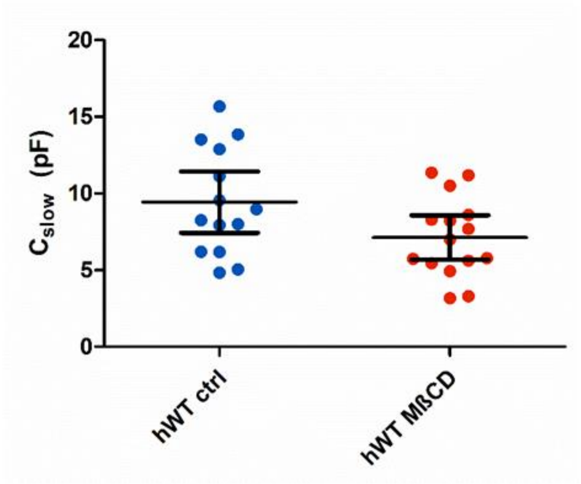


**Figure S10: Membrane capacitance.** Dot plot of the membrane capacitance ($C_{slow}$) for hWT ctrl and hWT MßCD. Error bars represent the 95% confidence interval. $C_{slow}$ gives us an indirect measure of the size of the cell. Both hWT ctrl and hWT MßCD have a similar distribution of cell sizes.

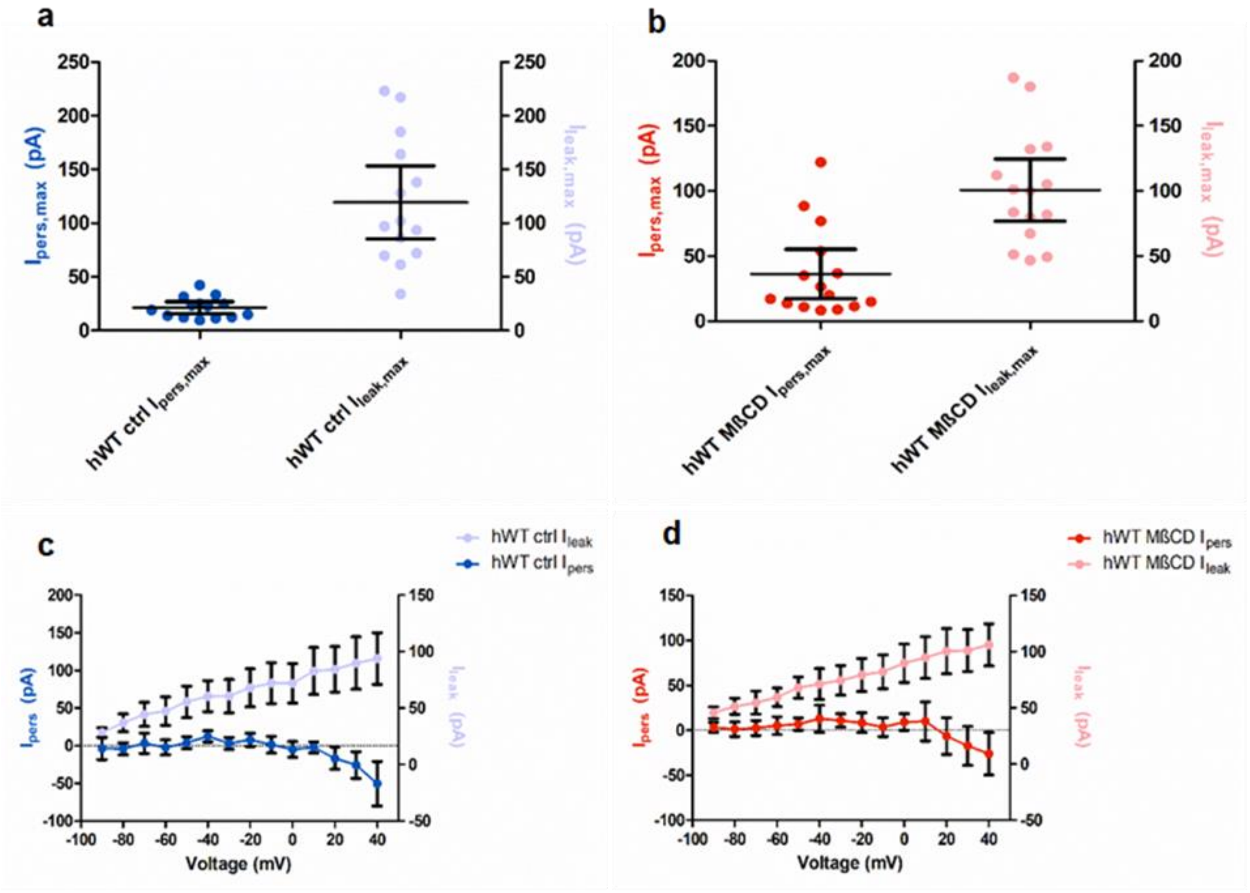

**Figure S11: A comparison of the persistent and leak currents. a.** Dot plot of the maximal persistent currents Ipeak,max (in pA) and maximal leak currents Ileak,max (in pA) for hWT ctrl. The left y-axis represents Ipers,max and right y-axis represents Ileak,max. Error bars represent the 95% confidence interval. **b.** Dot plot of the maximal persistent currents Ipeak,max (in pA) and maximal leak currents Ileak,max (in pA) for hWT MßCD. The left y-axis represents Ipers,max and right y-axis represents Ileak,max.Error bars represent the 95% confidence interval. **c.** Persistent currents and leak currents (in pA) vs voltage (mV) for hWT ctrl. The left y-axis represents Ipers,max and right y-axis represents Ileak,max. Error bars represent the 95% confidence interval. The magnitude of persistent currents are clearly much lower than that of the leak currents for every voltage step. **d.** Persistent currents and leak currents (in pA) vs voltage (mV) for hWT MßCD. The left y-axis represents Ipers,max and right y-axis represents Ileak,max. Error bars represent the 95% confidence interval. The magnitude of persistent currents are clearly much lower than that of the leak currents for every voltage step.

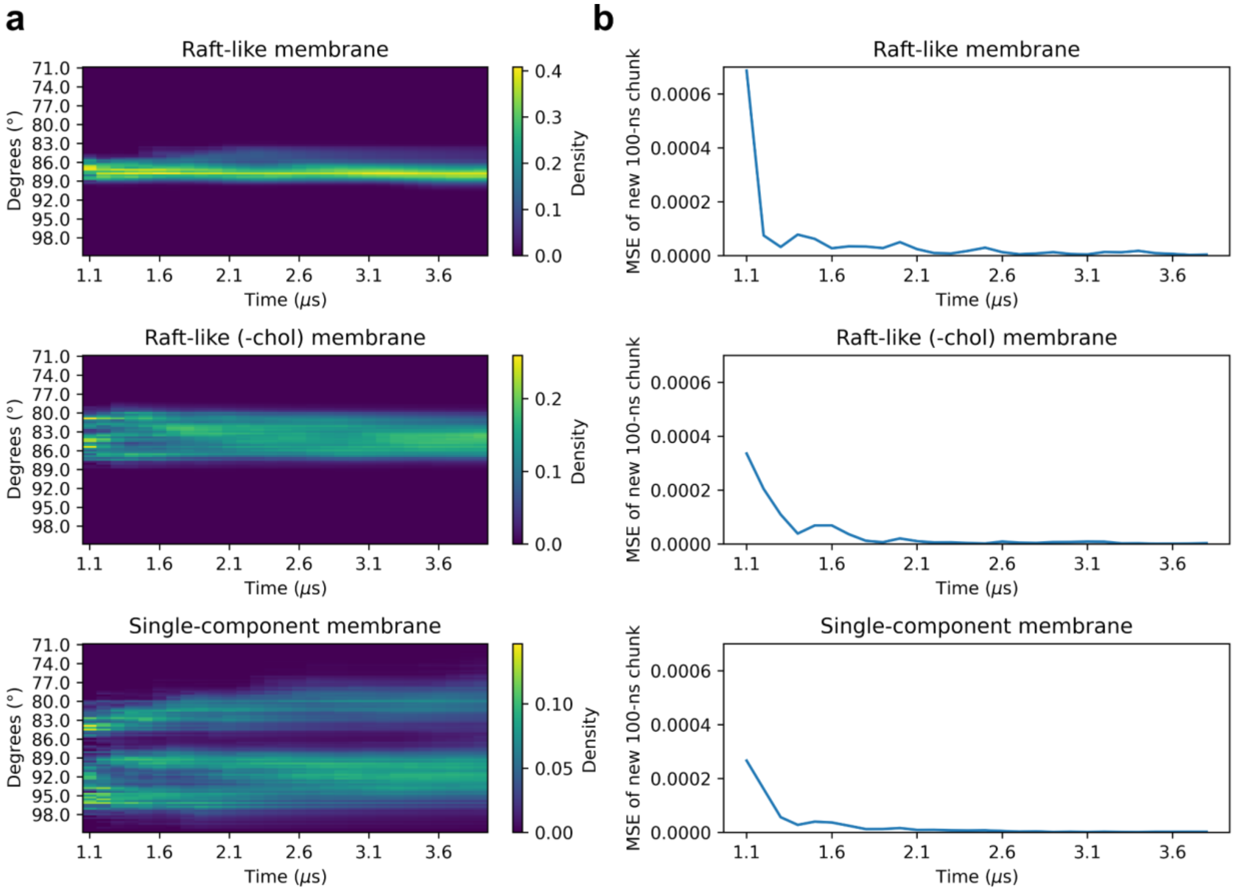


**Figure S12: Variation in the 4-fold symmetry of the channel over time**. Analysis of the temporal dynamics of the angle formed by the vectors connecting $VSD_{I}$ to $VSD_{III}$ and $VSD_{II}$ to $VSD_{IV}$. This angle is sampled every nanosecond across three replicas within each of the three membrane compositions. Starting from 1 microsecond, the distribution of these angle values is computed every 100 nanoseconds for each set of simulations corresponding to a particular membrane composition. (**a**) The evolving distributions are visualized as a heatmap, where each 100 ns interval updates the latest distribution. (**b**) Additionally, the figure quantifies the variation over time by calculating the Mean Squared Error (MSE) between successive distributions, providing insights into the dynamic changes in angle distribution within the system.

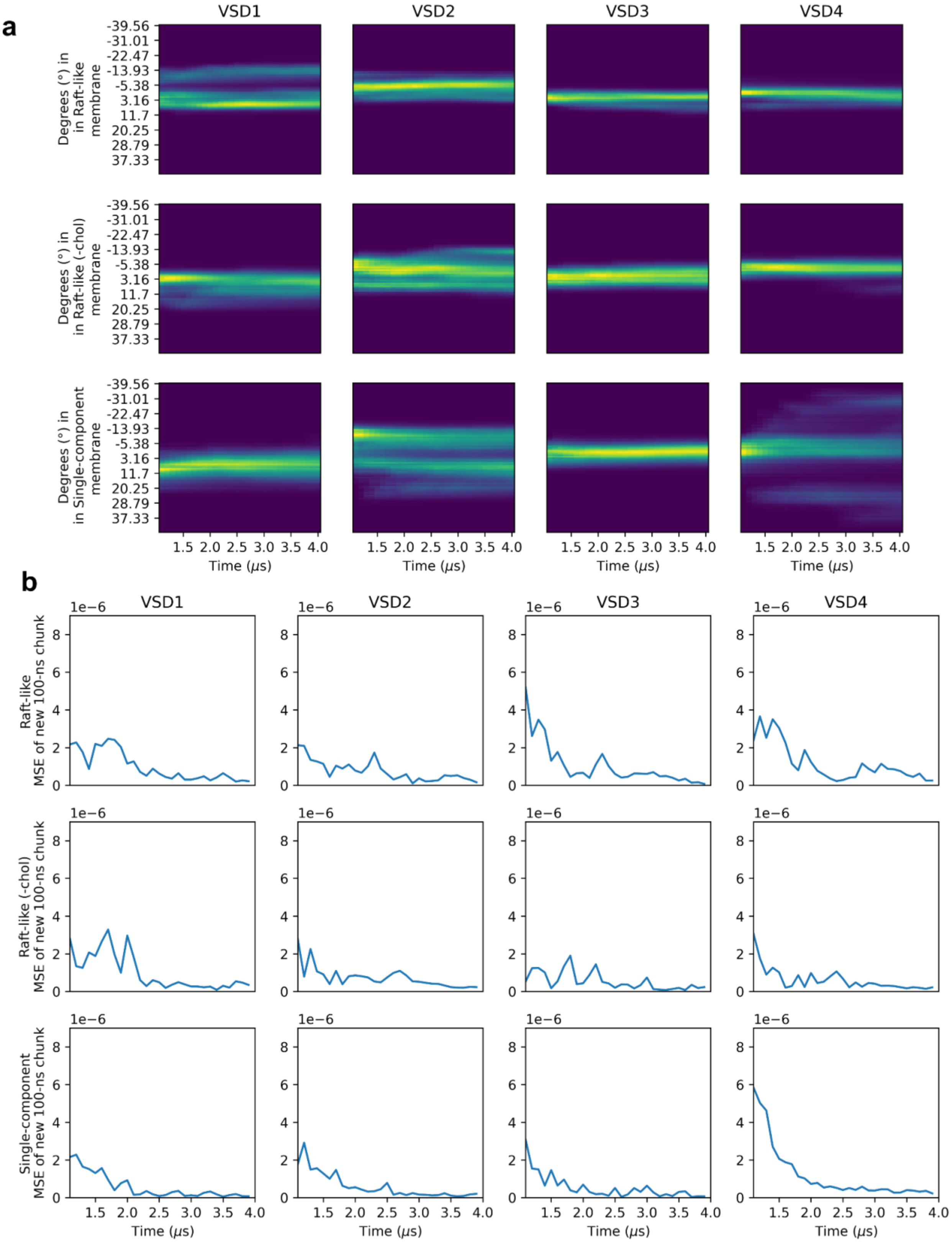
a
VSD1
VSD2
VSD3
VSD4
Degrees (°) in in Raft-like membrane
Degrees (°) in in Raft-like (-chol) membrane
Degrees (°) in in Single-component membrane
-39.56
-31.01
-22.47
-13.93
-5.38
3.16
11.7
20.25
28.79
37.33
1.5 2.0 2.5 3.0 3.5 4.0
Time (μs)
b
VSD1
VSD2
VSD3
VSD4
Raft-like MSE of new 100-ns chunk
Raft-like (-chol) MSE of new 100-ns chunk
Single-component MSE of new 100-ns chunk
1e−6
0
2
4
6
8
1.5 2.0 2.5 3.0 3.5 4.0
Time (μs)

**Figure S13: Variation in VSD Orientation Over Time.** Temporal evolution of the orientation of voltage-sensing domains with respect to the pore domain. The angle indicates which face of the VSD is oriented towards the pore domain, where an angle of 0 degrees corresponds to the state of the channel at t=0 μs. This orientation angle is sampled every nanosecond across three replicas within each of the three different membrane compositions, and for each VSD. Commencing from 1 μs, the orientation angle's distribution is computed at every 100-nanosecond interval for each simulation batch, aligned with a specific membrane composition. (**a**) The progression of these orientations is depicted as a heatmap, where the distribution is updated at each 100-nanosecond mark. (**b**) Furthermore, the figure quantitatively represents temporal variations by calculating the Mean Squared Error (MSE) between consecutive orientation distributions, offering a detailed view of the VSDs' dynamic orientation changes over time.
